# End-to-end proteogenomics for discovery of cryptic and non-canonical cancer proteoforms using long-read transcriptomics and multi-dimensional proteomics

**DOI:** 10.1101/2025.08.23.671943

**Authors:** Katarzyna Kulej, Asher Preska Steinberg, Jinxin Zhang, Gabriella Casalena, Eli Havasov, Sohrab P. Shah, Andrew McPherson, Alex Kentsis

## Abstract

Modern mass spectrometry now permits genome-scale quantitative measurements of biological proteomes. However, current approaches remain hindered by incomplete representation of biological variability of protein sequences in canonical reference proteomes, limiting studies to already annotated proteoforms. This, combined with reliance on tryptic peptides and limited sample complexity reduction techniques render detection of variant proteoforms missing from current reference annotations virtually impossible. Here, we report a method for end-to-end proteogenomics which integrates (i) sample-specific transcriptome assembly using long-read sequencing with (ii) multi-protease, multi-dimensional (MPMD) trapped ion mobility spectrometry (TIMS) parallel accumulation-serial fragmentation (PASEF) mass spectrometry. Termed ProteomeGenerator3, we benchmark its sensitivity and specificity for unbiased human cancer proteome discovery. Using long-read transcriptome sequencing of human Ewing sarcoma cancer cells, we detected over 40,000 full-length transcript isoforms, including aberrantly expressed neogenes, splice isoforms, and gene fusions. Combined with MPMD-TIMS-PASEF mass spectrometry, our proteogenomics workflow enabled the detection of 12,189 unique proteoforms with an average sequence coverage exceeding 50%, including 3,216 putative, non-canonical proteoforms, constituting one of the most complete human proteomes to date. Likewise, we report the proteogenomic performance metrics for current state-of-the-art implementations of DIA-NN, Spectronaut and diaTracer-MSFragger algorithms for data-independent acquisition (DIA) and PASEF mass spectral data, including optimal Double Rainbow-diaPASEF parameters for non-tryptic proteomes. In all, end-to-end integrative ProteomeGenerator3 provides a versatile platform for deep sample-specific proteogenomic discovery, suitable for the identification of disease biomarkers and therapeutic targets, and is available open-source from https://github.com/kentsislab/proteomegenerator3.

## INTRODUCTION

Recent advances in cancer genomics, transcriptomics, and proteomics have revealed extensive dysregulation of gene expression in cancer cells, driven by somatic genome rearrangements, aberrant transcriptional activation, alternative splicing, and post-translational modifications^1–4^. These alterations often result in the expression of novel or modified protein isoforms, neomorphic proteoforms, not catalogued in canonical reference databases^5–10^. Such proteins, including tumor-specific antigens and oncogenic drivers, can have direct roles in tumorigenesis and therapy resistance. Growing evidence underscores that the cancer proteome is significantly more heterogeneous than previously recognized, shaped by disease-specific events such as alternative transcription start sites, exon skipping, intron retention, RNA editing, and alternative translation reading frames^11–13^. These mechanisms give rise to both shared and patient-specific neomorphic proteoforms, including immune neoantigens, which have emerged as promising biomarkers and therapeutic targets. However, detecting these non-canonical proteoforms remains a major analytical challenge.

Mass spectrometry (MS)-based proteomics enables high-throughput, quantitative analysis of proteins^14–18^, yet its conventional bottom-up workflows depend on enzymatic digestion and peptide-spectrum matching (PSM) against reference databases^19–22^. This approach is limited in its ability to detect unannotated or sample-specific sequences, an issue especially pronounced in cancer, where the expressed proteome often diverges significantly from reference annotations. The completeness of the search database directly influences identification sensitivity, where sequence variants introduced by somatic mutations, splicing, or gene fusions are frequently missed.

Proteogenomic approaches address this limitation by integrating the analysis of genomic, transcriptomic and proteomic data to generate customized mass spectral analysis databases tailored to each sample^23–28^. These databases more accurately reflect the expressed proteome and improve the detection of tumor-specific proteins. Initially developed to support gene annotation, proteogenomics has matured into a key strategy in cancer research, aiding in the discovery of novel cancer drivers, neoantigens, and potential therapeutic targets^28–40^. Despite these advances, methodological challenges persist. Generating accurate proteogenomic databases from short-read RNA sequencing methods (RNAseq) can be hampered by difficulties in resolving transcript isoforms and identifying alternative open reading frames^23, 24, 26^. Simultaneously, trypsin-based protein digestion restricts sequence coverage, as certain proteins and variants yield tryptic peptides that are either suboptimal for MS detection or indistinguishable between similar proteoforms^41, 42^.

To overcome these obstacles, recent studies have embraced multidimensional proteomics workflows incorporating alternative proteases and orthogonal peptide fractionation^43–53^. Techniques such as high-pH reversed-phase (HpH-RP) chromatography, when combined with orthogonal separations, have been shown to improve protein identification and sequence coverage^54^. Top-down proteomics, which analyzes intact proteins, offers proteoform-level resolution but is constrained by limitations in resolving high-mass or low-abundance proteins and remains difficult to apply at scale^55^.

By integrating enhanced separation techniques, diverse proteolysis strategies, and integrative sample-specific MS analysis, bottom-up proteogenomic workflows can more effectively detect and quantify tumor-specific proteoforms. These advances are essential for comprehensive cancer proteome profiling and for enabling precision oncology approaches that target the full complexity of protein-level alterations in tumors, among many other biological applications. By combining sample-specific transcriptomes with mass spectrometry proteomics, we and others have demonstrated that an integrative proteogenomics framework can allow for enhanced detection of non-canonical sequences and peptides^56 57, 58^.

In parallel to proteogenomics methods development, long-read sequencing technology has been rapidly maturing^59, 60^, and several computational tools have been introduced to perform full-length transcript isoform analysis^61^. Benchmarking of long-versus short-read RNAseq in cancer cell lines has shown that long-read sequencing enables the elucidation of full-length, complex splice and fusion isoforms, both of which pose challenges for short-read RNAseq^62, 63^. To leverage these advances, we developed an approach to synthetically integrate these improved sequencing and proteomics methods (**Figure 1A**). To process long-read RNAseq data, we developed a new computational pipeline, ProteomeGenerator3 (PG3), which builds on the existing ProteomeGenerator (PG) methods^56, 57^ to generate sample-specific protein search databases from long-read RNAseq using (i) guided, de novo transcript assembly of full-length transcript isoforms and (ii) detection of fusion transcripts. In parallel, we acquired matched high-resolution mass spectrometry proteomics data for the same samples by performing multi-protease digestion combined with multi-dimensional peptide separation.

**Figure 1.**
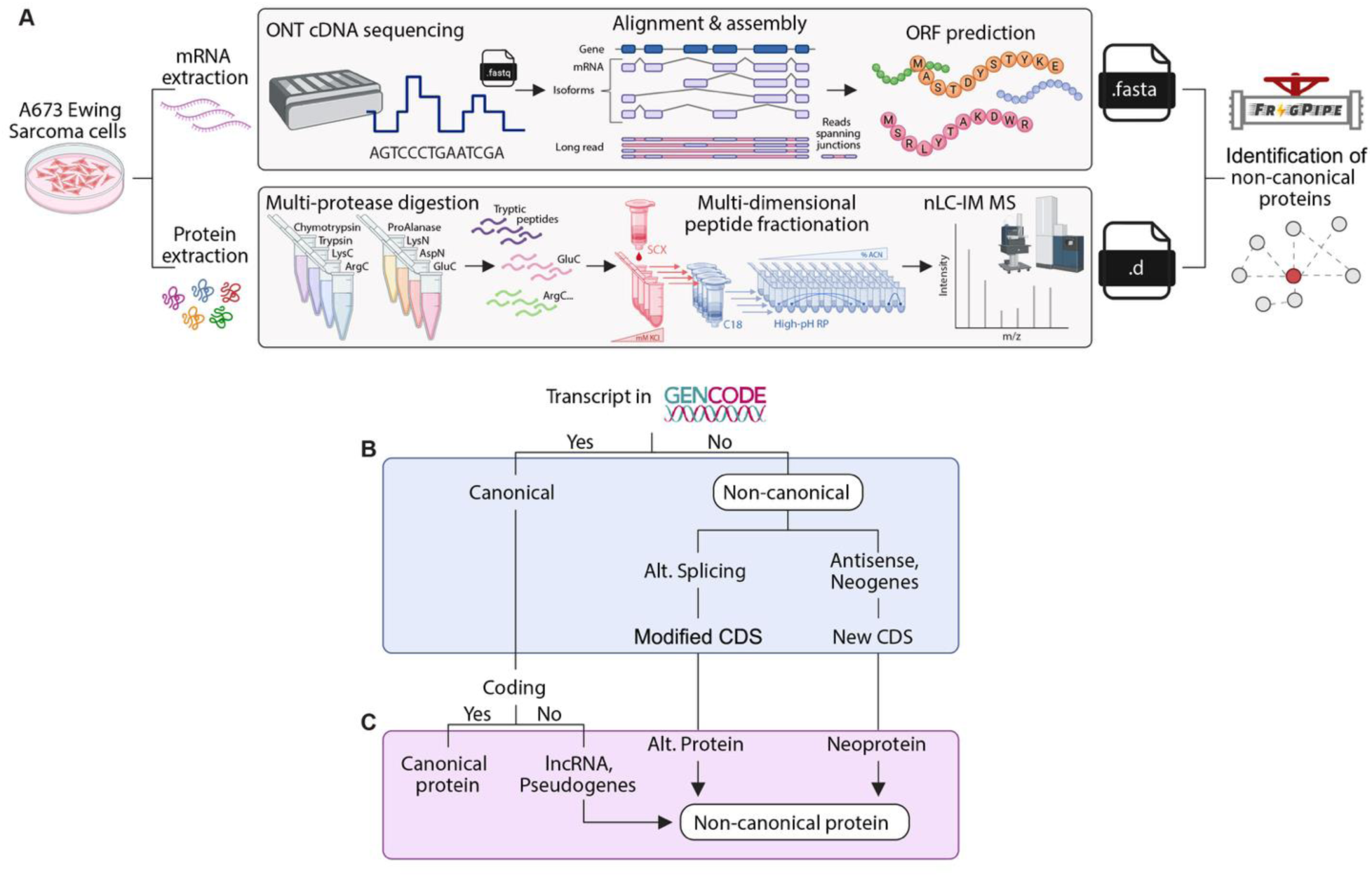
Schematic of integrated multi-dimensional proteogenomics framework for mapping tumor-specific proteoforms. **(A)** ProteomeGenerator3 workflow overview - transcriptomes and proteomes derived from the same biological sample are analyzed in parallel using high-coverage Oxford Nanopore Technologies (ONT) cDNA sequencing and high-resolution, high-accuracy multi-dimensional mass spectrometry. ProteomeGenerator3 processes the FASTQ-formatted mRNA sequencing reads to assemble predicted transcripts, determine coding sequences and splice isoforms, and generate FASTA-formatted proteogenomic databases. These databases include both canonical and non-canonical protein isoforms and are used in subsequent mass spectrometry searches. The resulting FASTA and diaPASEF-acquired timsTOF.d files are searched using FragPipe, enabling sensitive and comprehensive identification of expressed protein isoforms from the sample-specific proteogenomic database. **(B)** Transcripts from long-read sequencing originating coding sequences (CDS) are classified as “canonical” (annotated in GENCODE), and “non-canonical” (not annotated in GENCODE, including neogenes and alternative splicing isoform). **(C)** Sample-specific transcriptomes are used to search high resolution mass spectrometry data from orthogonally fractionated, multi-enzyme protein digests, allowing detection of non-canonical proteins, deriving from new transcripts or from known transcripts previously annotated as “non-coding”. *Created with BioRender.com;* https://BioRender.com/owf0tlx.

To further enhance sensitivity and improve the separation of singly and multiply charged as well as co-eluting peptides, we implemented parallel accumulation-serial fragmentation combined with data-independent acquisition (diaPASEF) on a trapped ion mobility spectrometry (TIMS) mass spectrometer coupled to time-of-flight (TOF) mass spectrometry^64^. This approach enables improved proteome coverage and quantitative accuracy across a broad range of sample loads. We then performed integrative computational analysis of these proteomics data using the sample-specific protein search database generated from long-read RNAseq transcriptome assemblies. We conducted the majority of these computational analyses using diaTracer-MSFragger^65, 66^, however, the PG3 approach is compatible with other mass spectral analysis algorithms including DIA-NN^67^ and Spectronaut^68^. This streamlined compatibility enables efficient processing of complex datasets, from peptide-spectrum matching to quantitative analysis, within a unified analytical framework.

To test the hypothesis that the PG3 approach can enable sensitive and specific detection of non-canonical proteoforms and neomorphic proteins (**Figure 1B-C**), we applied this method to human Ewing sarcoma A673 cells, which express the oncogenic EWSR1:FLI1 fusion transcription factor, known to regulate transcription via aberrant DNA binding, interaction with other transcription factors, and epigenetic reprogramming^69^. This chimeric factor can activate transcription from otherwise silent genomic regions, leading to the production of novel, spliced, and polyadenylated transcripts, known as neogenes, that are absent from standard reference databases from healthy human tissues and thought to be tumor-specific^69^. Ribosome profiling and proteomic data have confirmed that some of these transcripts are translated into tumor-specific proteins, providing a unique system for testing the sensitivity and specificity of improved proteogenomic tools. Thus, we sought to assess whether PG3 could capture the protein products of Ewing sarcoma neogenes and to explore the potential discovery of additional, previously unannotated transcriptomic and proteomic elements in human cells. As a result, we demonstrate that application of end-to-end integrative ProteomeGenerator3 improves our ability to study protein diversity in complex samples, enabling discovery of biological processes in physiological and pathological contexts.

## RESULTS

### ProteomeGenerator3: An end-to-end proteogenomics workflow for combining long-read transcriptomics with high-resolution mass-spectrometry proteomics

We developed a workflow for end-to-end proteogenomics (**Figure 2**) which combined long-read transcriptomics with multidimensional, mass spectrometry proteomics to allow for enhanced discovery of non-canonical proteins. We performed long-read complementary DNA sequencing (cDNAseq) generated from RNA isolated from the A673 cell line using the Oxford Nanopore Technologies (ONT) PromethION instrument; referred to as long-read/ONT RNAseq throughout the manuscript. To process the long-read RNAseq data, we used the nf-core/nanoseq pipeline (**Figure 2**, **Box 1**)^70^, performing quality control of raw reads with NanoPlot^71^ and FastQC^72^ and then aligning the sequencing reads to the GRCh38 (hg38) human genome reference using minimap2^73^. We obtained 5.2 million reads with a median Phred Score of 15 and median read length of 1.1 kB (**Supplementary Figure 1**). In parallel, JAFFAL was used to detect gene fusions by aligning reads to the human genome reference, identifying those which span multiple genes, and clustering these reads into breakpoint positions at exon boundaries^74^. After QC and alignment with the nf-core/nanoseq pipeline, the PG3 pipeline (**Figure 2**, **Box 2**) performs transcript assembly with Bambu^75^, which relies on probabilistic modeling to perform de novo transcript assembly and quantification. The probabilistic modeling allows users to fine tune the degree of false discovery acceptable for their analyses using the Novel Discovery Rate (NDR); NDR is an upper limit for the false discovery rate (FDR). We used Bambu to filter the data based on both the NDR and the number of full-length reads supporting a given transcript isoform. After assembly, quantification, and filtering, we used Transdecoder to predict ORFs for both transcript isoforms and gene fusions, where cDNA sequences are extracted from the transcript assembly using gffread. These predicted ORFs then become the protein search database for subsequent computational mass spectrometric analysis described below.

**Figure 2.**
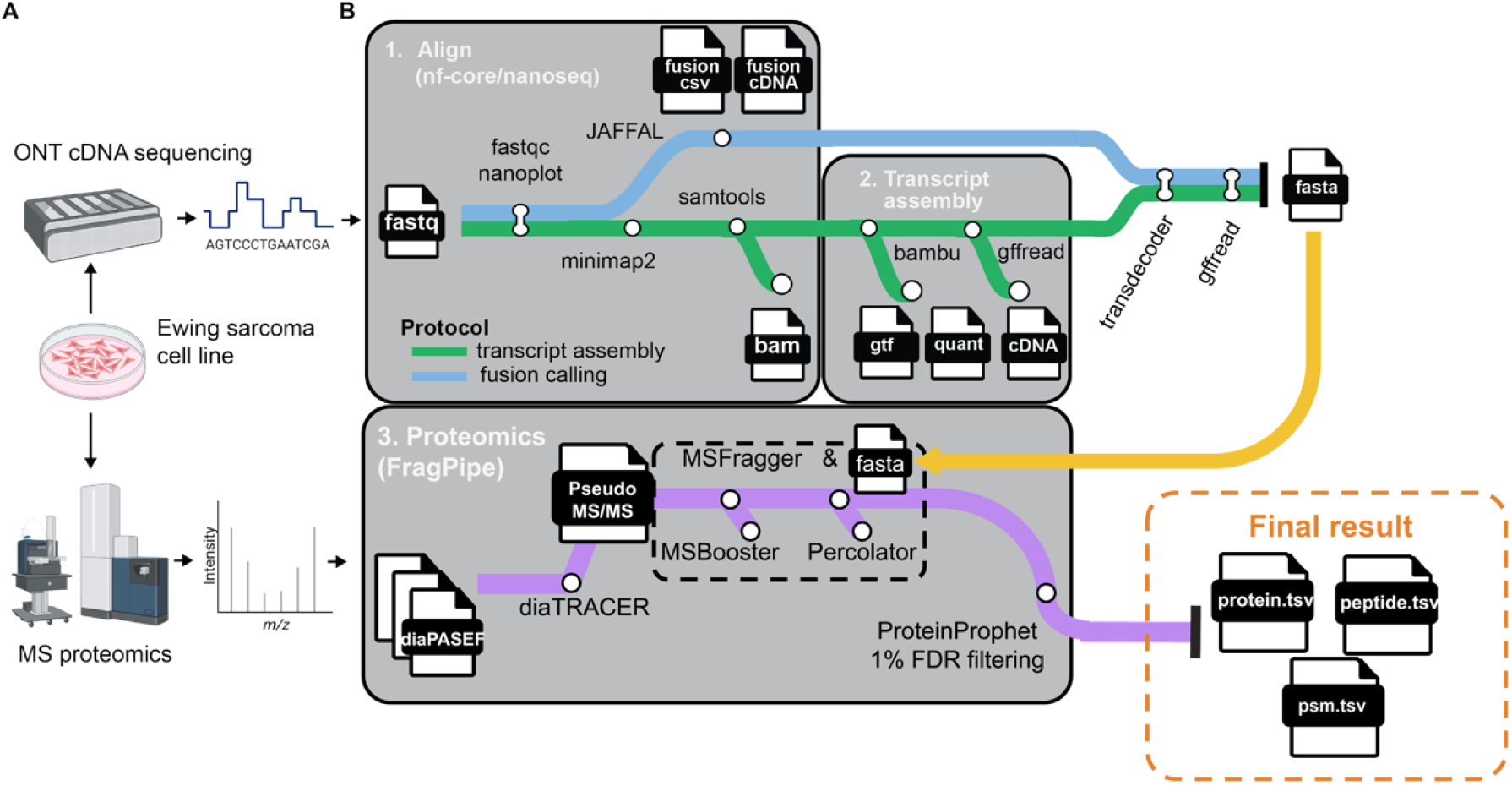
Overview of proteogenomics protocol. Oxford nanopore technologies (ONT) cDNA sequencing and MS proteomics were performed on the Ewing sarcoma cell line. Module 1 (labeled “Align”) shows alignment and quality control (QC) of long-read cDNA sequencing data using the nf-core/nanoseq pipeline and fusion calling with JAFFAL. Module 2 (labeled “transcript assembly”) shows transcript assembly performed with Bambu and extraction of cDNA sequences using gffread. Module 3 (labeled “Proteomics”) shows a simplified version of the Fragpipe diaTracer workflow which is utilized for peptide identification and quantification from diaPASEF-acquired data. diaTracer processes diaPASEF .d files to generate pseudo-MS/MS spectra (mzML) for MSFragger, enabling DDA-like database searching. This is followed by peptide-spectrum matching (PSM) using MSFragger, which is then rescored with deep learning-based algorithms via MSBooster and Percolator. Protein inference is conducted using ProteinProphet, and FDR filtering is applied at 1% FDR for PSM, ion, peptide, and protein levels using Philosopher. The Spectral libraries are generated directly from the DIA data using EasyPQP. The quantification is extracted from the DIA data using DIA-NN. *Panel A was created with BioRender.com;* https://BioRender.com/ctum5o6.

### Developing a multi-protease, multi-dimensional mass spectrometry proteomics strategy to maximize proteome sequence coverage

As many non-canonical proteins could be expressed at lower levels relative to canonical proteins, we first sought to develop a strategy to optimize the protein sequence coverage of the human proteome with our mass spectrometry proteomics. The proteome of the same A673 cell population from which we obtained long-read RNAseq data was analyzed using trapped ion mobility spectrometry-parallel accumulation-serial fragmentation (TIMS-PASEF) mass spectrometry, coupled to high-resolution, multi-protease multi-dimensional (MPMD) liquid chromatography. This integrated approach was used to enhance the detection of low-abundance and rare peptides, whose precursor selection for fragmentation is typically constrained by the limited duty cycle of conventional mass spectrometers. To maximize proteome sequence coverage of A673 cells using diaPASEF, we employed parallel digestion with eight sequence-specific proteases: trypsin, chymotrypsin, LysC, LysN, ArgC, GluC, AspN, and ProAlanase. These enzymes recognize distinct amino acid residues, enabling complementary cleavage patterns that increase peptide diversity and reduce sequence bias. This strategy enhances protein sequence coverage and supports the detection of non-canonical and low-abundance isoforms, which is especially important in complex samples. Furthermore, we performed sequential peptide fractionation by first applying strong cation exchange (SCX) chromatography to separate the sample into four fractions. Each SCX fraction was then further separated into 10 sub-fractions using high-pH reversed-phase (HpH-RP) chromatography^76–79^ (**Figure 3A**). Selected HpH-RP fractions, including flow-through, were concatenated, resulting in a total of 32 fractions per sample (**Figure. 3A**). This two-dimensional chromatographic fractionation approach takes advantage of the orthogonality between SCX and HpH-RP separation methods, which improves overall peptide separation prior to online low-pH reversed-phase LC-MS analysis ^78, 80^. Each fraction was individually analyzed by mass spectrometry, and the resulting data were then processed using the diaTracer and MSFragger algorithms within the FragPipe computational proteomics platform (**Figure 2**, **Box 3**)^65, 66^.

**Figure 3.**
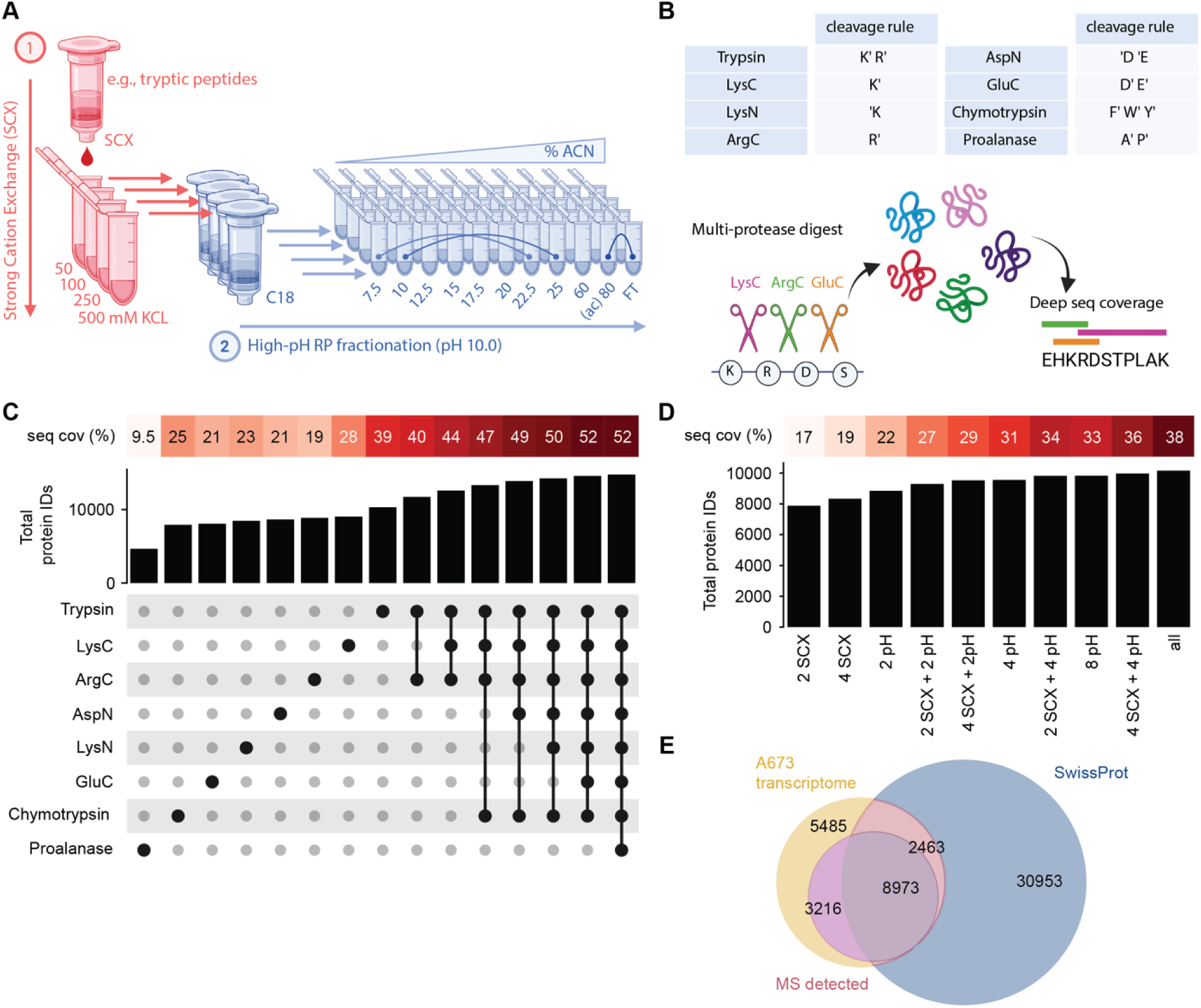
Quantification of protein IDs and sequence coverage of A673 proteome using multidimensional proteogenomics. **(A)** Schematic representation of the fractionation strategy employed in this study. A673 peptides were first fractionated using strong cation exchange (SCX) chromatography, with stepwise elution performed at increasing KCl concentrations of 50 mM, 100 mM, 250 mM, and 500 mM. Each SCX fraction was subsequently subjected to high-pH reverse-phase (RP) fractionation (pH 10). Peptides were eluted in 10 stepwise fractions with increasing acetonitrile (ACN) concentrations using a spin column. Selected fractions were concatenated into 8 final fractions, resulting in a total of 32 fractions per enzyme, which were then analyzed by LC-MS/MS. *Abbr. FT-flow-through.* **(B)** Schematic depicting the multi-protease digestion strategy that was employed for deep sequence coverage of proteins. The table summarizes all proteolytic enzymes used in this study along with their specific cleavage sites. Cleavage positions are denoted by a prime symbol (′), indicating whether the bond is cleaved on the N-terminal or C-terminal side of the specified amino acid residue. **(C)** Bar plot highlighting the total number of protein IDs identified with different enzyme combinations. Corresponding mean amino acid sequence coverage is shown above each bar. Matrix details which enzyme is used in the combination. **(D)** Bar plot illustrating the evaluation of different combinations of orthogonal peptide fractionation methods and/or varying numbers of peptide fractions to identify the most effective strategy for maximizing protein identifications and coverage. **(E)** Venn diagram showing counts of ORFs from the A673 transcriptome (A673 transcriptome, yellow), counts of proteoforms detected when diaPASEF proteomics data of the A673 cell line was searched using the A673-specific protein search database generated from the A673 transcriptome (MS detected, pink), and counts of all SwissProt human reference proteome (SwissProt, blue). *Panel A and B created with BioRender.com*; https://BioRender.com/2i48l39.

To evaluate if the MPMD proteomics approach improves the total number of protein IDs and depth of our sequence coverage, we assessed these metrics for different combinations of enzymes and fractions. The inclusion of proteolytic enzymes other than trypsin resulted in both a substantial improvement in the total number of proteins identified (42% increase) and in mean amino acid sequence coverage, with more than 10% increase compared to trypsin alone (52% vs. 39%, respectively) (**Figure 3B, C**). Gene Ontology (GO) term analysis of the identified proteins **(Supplementary Figure 2)** showed that, as expected, trypsin alone yielded the highest number of protein identifications and sequence coverage across cellular compartments, while different enzyme combinations enhanced coverage in a cell compartment-specific manner. For example, ArgC and LysC enhanced the number of IDs and sequence coverage for membrane proteins (**Supplementary Figure 2A**), whereas Trypsin, LysC and LysN provided the most IDs for nuclear proteins (**Supplementary Figure 2C**). We then aimed to establish the optimal combination of fractions to enable a more streamlined MPMD workflow without compromising proteome coverage **(Supplementary Table 1)**. We found that the HpH-RP fractionation appeared to have a greater impact on sequence coverage as compared to SCX; with four HpH-RP fractions for the same SCX fraction, we achieved an average sequence coverage of 31%, whereas four SCX fractions for the same HpH-RP fraction had an average coverage of 19% (**Figure 3A, D**). Overall, HpH-RP fractionation accounted for the majority of protein identifications and sequence coverage when combined with SCX, highlighting its complementary role in enhancing proteome depth using the TIMS-diaPASEF approach.

In typical DIA mass spectrometry methods, TIMS isolation parameters are optimized for tryptic peptides, which predominantly yield multiply charged precursors within a well-characterized ion mobility (1/K₀) and m/z space. However, alternativ e proteases such as GluC, AspN, or chymotrypsin generate peptides with distinct physicochemical properties, including shorter sequences and a higher prevalence of singly charged ions. These non-tryptic peptides often fall outside standard ion mobility and m/z isolation windows, leading to suboptimal sampling and reduced identification rates.

To overcome this, we first analyzed chymotryptic digests using a standard DDA-PASEF method. As expected, the default TIMS isolation polygon, designed for multiply charged tryptic peptides, excluded a significant portion of the singly charged ion population, which appeared as a distinct cluster outside the main precursor cloud (**Supplementary Figure 3A**). This approach yielded approximately 18,574 unique peptides per injection of an unfractionated chymotryptic A673 proteome digest, with the majority (∼77%) being doubly charged (**Supplementary Figure 3E, I**). Singly charged ions were largely excluded, contributing to only 0.01% of identified peptides (**Supplementary Figure 3E**).

To improve peptide ion precursor coverage, we modified the standard DDA-PASEF acquisition in two ways: by removing the default TIMS isolation polygon (**Supplementary Figure 5B**) or by extending the precursor ion mobility range from 1.6 to 1.7 1/K₀ (**Supplementary Figures 3C**). These modifications enabled the fragmentation of singly charged precursors, which accounted for approximately 34% and 38% of all identified peptides per run in the non-polygon and extended-polygon methods, respectively (**Supplementary Figures 3F, G**). Additionally, we tested a previously reported acquisition strategy, Thunder DDA-PASEF, which specifically targets the singly charged ion region (**Supplementary Figure 3D**)^81^. This approach further increased the identification of singly charged peptides to ∼45%, though it came at the expense of reduced identification of multiply charged (≥+2) peptides (**Supplementary Figures 3H**). The total counts of peptides and proteins identified by each method are summarized in **Supplementary Figure 3J–K**.

Encouraged by the performance of the Thunder DDA-PASEF approach, we next implemented its improved DIA derivative, termed Double Rainbow-diaPASEF, to enable comprehensive sampling of both singly and multiply charged precursors under DIA acquisition (**Supplementary Figure 4A, C**). Compared to the standard polygon, the double rainbow isolation scheme captured a higher proportion of +1 and +2 peptide precursor ions (**Supplementary Figure 4B-D**), leading to an increased identification of short peptides, particularly those ranging from 7 to 13 amino acids (**Supplementary Figures 4E-G**). In contrast, the standard TIMS polygon more effectively targeted precursors with charge states of +3 or greater. Notably, the broader precursor coverage of the Double Rainbow-diaPASEF method resulted in ∼40% fewer MS2 scans per cycle compared to the standard DIA method (**Supplementary Figure 4H**), which had implications for the overall number of peptide and protein identifications. Despite this, Double Rainbow-diaPASEF enabled the identification of 2,380 unique peptides corresponding to 263 proteins not identified in the standard DIA method (**Supplementary Figure 4I**). Notably, gene set enrichment analysis (GSEA) showed enrichment of membrane-bound organelle proteins, consistent with the observation that chymotrypsin enhances sequence coverage of membrane proteins relative to trypsin (**Supplementary Figure 4J**)^54, 82, 83^. Together, these results demonstrate that optimized MS acquisition strategies can enhance the detection of peptides derived from non-tryptic or mixed enzymatic digestions, thereby improving overall proteome coverage.

### Sample-specific long-read transcriptomics reveals non-canonical transcript isoforms expressed in human cancer cells

Our next goal was to assess whether PG3 could detect previously reported transcripts and to investigate the presence of additional unannotated transcriptomic elements. We found that the A673 cell transcript assembly contained 40,038 transcript isoforms with >1 full-length reads; 23,274 of these isoforms had ORFs predicted by Transdecoder^84^ (**Figure 4A**). By comparing transcript structures, we found that 38,917 transcript isoforms matched to the GENCODE v45 human genome annotation, whereas 13,607 transcript isoforms matched those found in the long-read transcript assembly from 88 samples derived from human tissues and cell lines in the Genotype-Tissue Expression Project (GTEx v9) ^85^, which we assembled using PG3 for comparison. In the A673 transcript assembly, we observed 1,121 transcript isoforms which were not annotated in GENCODE v45, which we term “non-canonical” transcript isoforms (**Figure 4A**). Of these 1,121 non-canonical transcript isoforms, 123 matched to transcripts detected in GTEx tissues and cell lines. Many of these transcripts were expressed at various levels across all healthy GTEx tissues (**Supplementary Figure 5A**), with the highest expression in fibroblasts, brain, and lungs (**Supplementary Figure 5B**). Detected non-canonical transcript isoforms exhibited an increased frequency of alternative first exons and exon skipping (**Figure 2B**).

**Figure 4.**
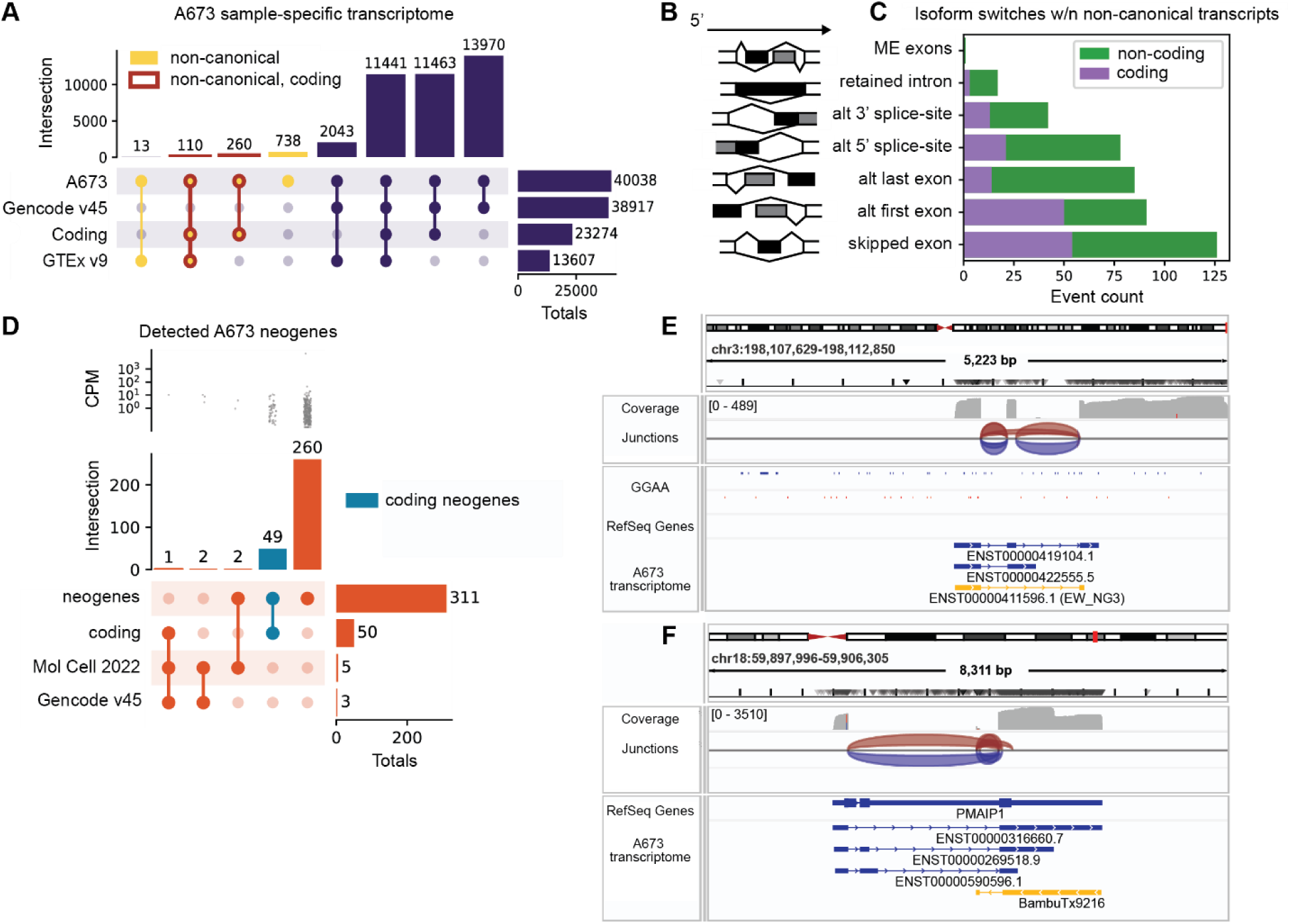
Long-read transcriptomics of Ewing sarcoma cell line detects non-canonical transcriptional events. **A**. Upset plot comparing long-read transcript assembly of the A673 cell line to GENCODE v45 and GTEx v9 ^85^. Vertical bars show intersection sizes and horizontal bars show total counts for each set within the A673 transcript assembly. “A673” row in the upset matrix refers to all transcripts detected from long-read transcript assembly of the A673 cell line. The remaining rows depict the subsets of A673 transcripts which overlapped with the Gencode v45 and GTEx v9 annotations, as well as what subset of A673 transcripts predicted to be coding by Transdecoder^84^. “Non-canonical” refers to transcripts not found in Gencode v45. **B-C**. Bar plot of alternative (alt) splicing events detected within non-canonical transcripts in the A673 cell line. Alt splicing events across all transcripts are shown in green whereas those found in transcripts with predicted ORFs are shown in purple. ME exons refers to mutually exclusive exons. **D**. Upset plot depicting putative neogenes detected in the A673 transcript assembly. Strip plot in top panel shows expression levels of neogenes quantified using Bambu ^75^ in counts per million (CPM). In the upset plot, vertical bars show intersection sizes of sets and horizontal bars show total counts for each set. The “Mol Cell 2022” row shows the neogenes initially detected in ^3^, which were also found in the Gencode v45 annotation. (**E**) IGV snapshot showing the detected neogene from ref.^3^ in yellow (labeled “ENST00000411596.1 (EW_NG3)”). The GGAA track shows microsatellite binding regions nearby to the transcriptional start site. (**F**) IGV snapshot of one of the A673-specific neogenes discovered in our long-read RNAseq (shown in yellow and labeled “BambuTx9216”).

Recent analyses of A673 cells found evidence of transcriptional activity in genomic regions which are normally inactive^3^. The transcriptions of these “neogenes” is driven by the EWSR1:FLI1 fusion protein acting as an oncogenic chimeric transcription factor which binds to microsatellite regions upstream of cryptic transcriptional start sites. We observed 327 transcripts which mapped to 311 genes that were not present in the GENCODE v45 annotation and could potentially represent neogenes (**Figure 4D-F**). We also confirmed the expression of five previously reported neogenes^3^; three of which matched to genes found in GENCODE v45. For these recovered neogenes^3^, we observed GGAA microsatellite sequences near their transcriptional start sites, consistent with the dependence on EWSR1:FLI1. Some of the newly discovered neogenes appeared in genomically active regions but were anti-sense to canonical transcripts with distinct sets of transcribed exons (**Figure 4F**). Additionally, we found 65 neogenes occurring in gene deserts when comparing to both GENCODE and RefSeq human genome annotations (**Supplementary Figure 6**). Thus, PG3 analysis of A673 Ewing sarcoma cells revealed extensive transcriptomic complexity, including numerous non-canonical and neogene transcript isoforms not captured by existing genome annotations.

### End-to-end proteogenomics allows for the discovery of cryptic and non-canonical proteoforms in human cancer cells

Encouraged by these findings, we used ORFs predicted from the A673 transcriptome assembly as a protein search database to detect non-canonical proteoforms. We used diaTracer combined with MSFragger to analyze matching DIA mass spectral data, as implemented in Fragpipe^65, 66^. We detected 12,189 proteoforms corresponding to 60% of the predicted A673-specific ORFs (**Figure 3E**). Based on sequence identity, 8,973 of the detected proteoforms matched canonical SwissProt human protein sequences, while 3,216 represented non-canonical proteoforms with no corresponding SwissProt matches (**Figure 3E**).

Given the widespread adoption of several different DIA mass spectral analysis algorithms, we sought to assess the sensitivity and specificity of diaTracer-MSFragger (v22.0), Spectronaut (v19.0), and DIA-NN (v2.1.0) for proteogenomic studies. To do this, we introduced 100 *Archaebacteria loki* sequences into the human protein search database to serve as false positive decoys (chosen by randomly sampling all *A. loki* sequences in SwissProt), taking advantage of their evolutionary distance and lack of sequence homology to human proteins^86, 87^. This allowed us to estimate the rate of false-positive proteogenomic identifications in a controlled manner. All three algorithms showed similar specificity, identifying approximately 1-2% *A. loki* peptides at a 1% FDR at both the peptide and protein levels (**Supplementary Figure 7A-B**). Moreover, the total number of protein identifications across the algorithms remained comparable, indicating their equivalent overall sensitivity for proteogenomic analyses (**Supplementary Figure 7A-B**).

To determine if the 3,216 tumor-specific proteoforms detected in the A673 cells that did not match currently annotated SwissProt proteoforms were bona fide non-canonical proteoforms, we assessed their apparent sequence similarity against the entire UniProt database using BLASTp (**Figure 3E**). This identified 72 putative non-canonical proteoforms with unique peptides which had no BLASTp matches to any UniProt proteoforms (**Supplementary Table 3**). Further, we ran BLASTp searches of the identified peptides themselves against both UniProt and NCBI non-redundant protein sequences and found that only two proteins had peptides which had matches in these databases. We found that 22 of these proteoforms were uniquely detected as tryptic peptides, however other enzymes also contributed to the discovery of these unknown proteoforms (**Figure 5A**). For example, the oncogenic EWSR1:FLI1 fusion protein itself was identified by a unique junction spanning peptide in the chymotrypsin digest (**Supplementary Figure 8**). To assess the likelihood that these proteoforms were not due to spectral mismatches, we compared posterior error probability (PEP) mass spectral identification scores for detected non-canonical proteoforms to the identified canonical SwissProt proteoforms (**Figure 5B**). We observed that a subset of the identified non-canonical proteoforms had PEP mass spectral scores equal to or better than the median PEP scores of canonical SwissProt proteoforms, supporting the accuracy and confidence of their identification (**Figure 5B**). To test the hypothesis that the spectra of these 72 putative non-canonical proteoforms are better matches to canonical peptides, we conducted a search against a protein search database constructed by concatenating the A673-specific preoteome with SwissProt, allowing spectra to directly compete for A673 non-canonical peptides and all possible canonical peptides. We found that 17 of the 72 putative non-canonical proteoforms still had spectral matches to novel peptides in the concatenated A673 + SwissProt search (**Supplementary Table 4**), supporting that these spectral matches are derived from non-canonical proteoforms.

**Figure 5.**
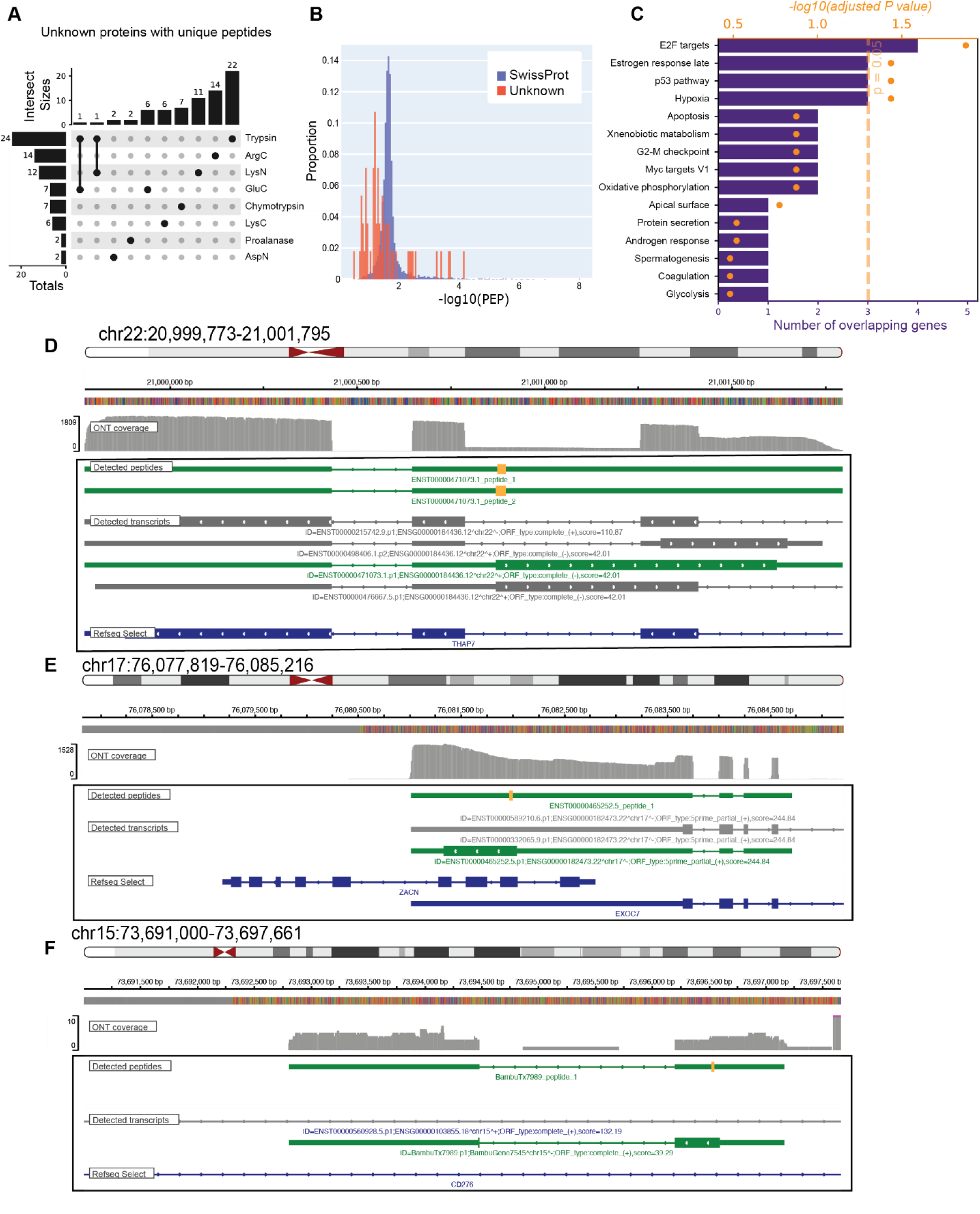
Identification of non-canonical proteins using end-to-end proteogenomics protocol. (**A**) Upset plot showing proteins with unique peptides from Figure 4A which had no BLASTp match to any protein in UniProt. (**B**) Histograms of posterior error probability (PEP) scores for SwissProt proteins and the “unknown” proteins from panel A, all of which were identified in the sample-specific A673 computational proteomics search with Fragpipe. PEP scores are the mean of all unique peptides for a given protein. **C)** Gene set enrichment analysis performed for the unknown proteins shown in panels *A-B* using the MSigDB Hallmark 2020 gene sets. (**D-F**) Examples of unknown proteins identified at the THAP7, EXOC7, and CD276 loci, respectively. For each panel, tracks from top to bottom show: ideogram, ONT read coverage, detected peptides (orange) shown on transcript structure (green), detected transcripts shown with predicted ORFs (the unknown protein is highlighted in green), and RefSeq Select.

The majority of detected non-canonical proteoforms originated from alternative open reading frames (ORFs) embedded within otherwise canonical transcript isoforms annotated in GENCODE. For these genes, we performed gene set enrichment analysis (**Figure 5C**) and found that several were related to transcription factors and cancer-related pathways (e.g., the p53 pathway and Myc Targets V1). At the *THAP7* locus, we found evidence of an anti-sense ORF relative to ENST00000471073.1, with two unique ArgC peptides (PEP = 2e-2) and Pfam structural domain match (**Figure 5D**)^88, 89^. At the *EXOC7* locus, we found a unique tryptic peptide (PEP = 2e-4) related to an alternative ORF relative to ENST00000465252.5 (**Figure 5E**), with evidence of Pfam membrane and signal peptide domains (**Supplementary Figure 9**). Lastly, in the proalanase digest, we found a unique peptide match (PEP = 6e-2; 1 Pfam domain match) associated with an anti-sense neotranscript within the intronic regions of *CD276* (B7-H3), a known cancer-associated protein, which is a current target for investigational cancer therapies (**Figure 5F)**. Overall, these results indicate that sample-specific integrative proteogenomic approach using ProteomeGenerator3 enables accurate and sensitive discovery of non-canonical disease-specific protein variants.

## DISCUSSION

Advances in clinical genomics have enabled progress in precision oncology, but new tools which leverage unbiased and sensitive analysis of disease and sample-specific proteoforms are necessary to elucidate complex tumor biology on the protein level. In the current work, we present an improved end-to-end proteogenomics approach for integration of long-read transcriptomics with high-resolution mass spectrometry proteomics which enables the discovery of non-canonical and neomorphic proteoforms. By using a sample-specific, long-read transcript assembly as the substrate for mass spectral analysis, we identified over 3,000 non-canonical proteoforms (**Figure 3E**), with 72 of these protein variants (**Figure 5A**) having no apparent matches to any protein isoforms annotated in current human proteome databases. Notably, multi-protease, multi-dimensional (MPMD) trapped ion mobility spectrometry (TIMS) parallel accumulation-serial fragmentation (PASEF) mass spectrometry enhanced protein sequence coverage depth to levels which cannot be achieved with a single enzyme and/or fraction (**Figure 3A-B**).

By testing different combinations of enzyme digests, we found that while trypsin alone had a sequence coverage of 39%, this could be boosted to 52% by inclusion of seven additional enzymes (**Figure 3C**). Trypsin and LysC alone had the highest coverage individually, which is consistent with prior studies^54^. Each of the 8 enzymes used in the study revealed non-canonical proteoforms with no homology to known proteins (**Figure 5A**), suggesting that multi-enzyme approach is necessary to capture non-canonical proteoforms. In terms of HpH-RP and SCX fractionation, HpH-RP appeared to have a stronger impact than SCX. In fact, doubling the number of pH fractions from 2 to 4 for the same SCX fraction led to an increase in sequence coverage from 22 to 31%. In contrast, doubling the number of SCX fractionations from 2 to 4 increased the coverage from 17 to 22% (**Figure 3D**). In addition to its contribution to proteome coverage, HpH-RP offers several practical advantages over SCX. HpH-RP fractionation typically provides higher sample recovery and minimizes sample loss, as it avoids the additional desalting steps often required before or after SCX-based fractionation. Consequently, fractionation strategies that provide more effective separation, generate cleaner fractions, and minimize sample handling and loss, such as HpH-RP, was considered optimal for streamlined proteomic workflows. These insights on the optimal combination of enzymes and fractionation should inform experimental design for the detection of non-canonical proteoforms in cancer and other biological contexts for which proteome annotation is incomplete.

In our workflow, we implemented FragPipe due to its open-source and modular architecture, which integrates MSFragger and diaTracer algorithms, offering the flexibility and control necessary for complex proteogenomics applications. MSFragger enables fast and scalable database searches, making it well-suited for large datasets, while diaTracer expands this functionality by supporting open (mass-offset) searches for the unbiased identification of PTMs from DIA-PASEF data. In contrast, Spectronaut and DIA-NN do not currently support native mass-offset searches using DIA-PASEF acquisitions, though they exhibit comparable proteogenomic performance to diaTracer-MSFragger.

To improve coverage of non-canonical and singly charged peptides, we optimized the TIMS selection polygons used in our DIA-PASEF acquisition scheme. While conventional methods prioritize doubly and triply charged tryptic peptides, our empirical mapping of chymotryptic samples revealed a substantial population of singly charged precursors outside standard acquisition windows. By refining the DIA-PASEF polygon design, guided by DDA-PASEF and supported by Thunder-PASEF findings^81^, we developed a customized dual-row scheme (Double Rainbow-diaPASEF) that significantly expands coverage of this precursor space, enabling improved detection of peptides arising from non-tryptic or poorly characterized enzymatic activity. Combining this tailored acquisition strategies with the standard polygon approach enhances the detection of peptides from non-tryptic or mixed enzymatic digestions, thereby increasing overall proteome coverage in both DDA and DIA mass spectrometry.

We computationally probed the hypothesis that the candidate non-canonical proteoforms are false MS identifications by performing a search against a protein search database composed of the A673-specific proteome and SwissProt, which yielded additional support for 17 proteoforms. However, future experimental validation will be necessary to validate and determine if the candidate non-canonical proteoforms identified in our study are functional, as well as the mechanisms that cause their expression. For example, non-canonical proteoforms directly regulated by the oncogenic fusion EWSR1:FLI1 would represent ideal tumor-specific biomarkers and therapeutic targets.

In all, this work lays the foundation for developing improved integrative methods with maximal sensitivity and specificity for the elucidation of proteomic diversity deriving from complex biological processes and those dysregulated in disease such as cancer. While previous studies have examined the use of proteogenomics-derived protein search databases and multi-enzyme/fractionation approaches separately, our work here shows that these insights can be combined to yield a highly sensitive approach using MPMD mass spectrometry. The combined PG3 MPMD approach we establish here should enable the discovery of non-canonical proteoforms in various human disease states and other organisms for which genome, transcriptome, and proteome are not well annotated. Moreover, this approach is designed to take advantage of improvements in transcriptome sequencing coverage and MS techniques for peptide separation and digestion, thus the sensitivity of our method is poised to improve. This approach can enhance our understanding of the biological processes underlying diseases like cancer and facilitate the identification of potential cancer-specific therapeutic targets.

## METHODS

### Cell culture

Authenticated (STR resulted in a 98% match), mycoplasma negative A673 Ewing sarcoma cells were grown at 37 °C with 5% CO_2_ in DMEM medium (Thermo, Gibco) supplemented with 10% heat-inactivated fetal bovine serum (FBS), 1% L-glutamine, and 1% Penicillin/Streptomycin solution. Cells were collected using an Accutase (Thermo, Invitrogen) at >80% confluency through centrifugation at 1500 rpm for 5 min at room temperature (RT). The supernatant was removed, and cells were washed twice with PBS and centrifuged at 1500 rpm for 5 min at RT. The resulting cell pellet was stored at −80 °C.

### RNA extraction

Frozen cells were lysed in Buffer RLT with 2-mercaptoethanol and RNA was extracted using the RNeasy Mini Kit (Qiagen) on the QIAcube Connect (Qiagen) according to the manufacturer’s protocol. Samples were eluted in 55 µL RNase-free water.

### Long-read sequencing

After RiboGreen quantification and quality control by Agilent BioAnalyzer, 28.8 µg of total RNA with a RIN of 7.1 underwent cDNA synthesis and library preparation using the Ligation Sequencing Kit v14 (Oxford Nanopore) according to instructions provided by the manufacturer. mRNA selection was performed with a VN Primer (IDT) and RT and strand switching proceeded with Maxima H Minus Reverse Transcriptase (ThermoFisher) and a strand-switching primer (IDT) for 90 minutes at 42°C. After second strand synthesis, the NEBNext Ultra II End Repair/dA-tailing Module (New England Biolabs) was used to prepare cDNA for adapter ligation with buffer and adapters from the Ligation Sequencing Kit and NEBNext Quick T4 DNA Ligase (New England Biolabs). Samples were sequenced on a PromethION 24 using the R10.4.1 flow cell (Oxford Nanopore). 52 million reads were generated for the sample with an N50 of 1.9 kilobases. Basecalling was performed with Guppy (version 6.5.7) using the high-accuracy model, 400 bps.

### Transcript assembly and quantification of A673 cell line RNAseq data

As described in **Figure 2**, we used the nf-core/nanoseq pipeline version 3.0.0 (Ewels et al. 2020) to perform QC, align the long-read cDNA sequencing data, and perform fusion calling with JAFFAL. To assess read quality, nanoplot (v1.42.0) was used (output from nanoplot shown in Supplementary Figure 1). Minimap2 (v2.17-r941) was used with the flags “-ax splice” to align raw reads to the GRCh38.p14 primary genome assembly (Gencode release v45) and samtools (v1.15.1) was used to convert SAM file output to a coordinate-sorted BAM file. The aligned bam was then passed to the ProteomeGenerator3 pipeline, and transcript assembly and quantification was performed with Bambu (v3.5.1), which was run in several steps. First, transcript read classes were generated by running Bambu in discovery mode (to allow for use with future analyses), reference-guided, de novo transcript assembly was then first performed with a novel discovery rate (NDR) = 1, followed by filtering of this new transcriptome annotation for transcripts with NDR < 0.1. Bambu was then run in quantification mode with this annotation to determine read support, and the final transcript assembly was filtered for transcripts with > 1 full-length read (which has been demonstrated to improve specificity^75^.

### Analysis of transcript structure and alternative splicing patterns

The comparisons of the A673 transcript assembly to Gencode v45, GTEx v9^85^, and ref. ^3^ in **Figure 4** was performed using gffcompare with default settings. To compare to ref.^3^ we took Table S3 from the supplement of their manuscript, converted it to a gtf, then performed liftover from hg19 to hg38 using the UCSC LiftOver commandline tool (downloaded 06/24/2024) along with the hg19ToHg38.over.chain.gz file (the “-gff” flag was used as files were in gtf format). We then filtered the gtf for Ewing sarcoma specific neogenes (those which had transcript ids which had prefixes “Ew_NG”) and used gffcompare to compare to our A673 gtf. To analyze alternative splicing patterns (**Figure 4C**), SUPPA2 was used with the “--pool-genes” flag and the “-b V” flag for variable boundaries. For the expression data for overlapping transcripts between the A673 non-canonical transcriptome and GTEx v9 appearing in **Supplementary Figure 5**, the quantification data provided from ref.^85^ was used. To determine which of the neogenes in the A673 transcript assembly were in “gene deserts” (i.e., parts of the canonical annotation where no known genes exist), we used bedtools intersect to determine which transcripts in the A673 assembly do not overlap with any transcripts in RefSeq and Gencode (we first converted the A673 transcript assembly and RefSeq to bed files, and the Gencode v45 gtf to gff).

### ORF prediction and generation of protein sequence search database

To generate a transcript fasta file from the gtf output of Bambu, gffread was used. The Bambu transcript fasta file and the JAFFAL fusion fasta file were then fed to Transdecoder to predict ORFs with a minimum ORF length of 100 AAs and only allowing a single ORF per transcript (the “--single_best_only” flag). A custom python script was used to format these fasta files for use with Fragpipe and to take only complete ORFs.

### Protein sequence coverage calculation

The weighted mean protein sequence coverage (*C̄*_*wt*_) across multiple enzymes was calculated by first calculating the cumulative peptide length per protein (*L_pep_*). We define this as the number of amino acid (AA) positions which are covered by peptides for a given protein, accounting for overlap between the detected peptides. Then to compute *C̄*_*wt*_ for a given combination of enzymes we used the following equation:

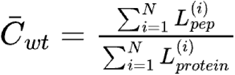

Where *N* is the total number of detected proteins and ***L**_protein_* is the length of a given protein *i*. For all figures the peptide.tsv output file from Philosopher^90^ was used to the determine which AA positions were covered, with the exception of **Figure 3D**, where MSstats^91^ output was used as it specifies which peptide was detected by different fractions.

### GO term annotation

To perform the analysis of protein sequence coverage across different cellular components as shown in **Supplementary Figure 5**, we used the biomaRt package to download GO terms for Ensembl v111. We then matched detected proteins using the Ensembl IDs of their corresponding transcripts. For the subpanels, we filtered for proteins which matched exclusively to “membrane”, “cytoplasm”, and “nucleus”, using their cellular component annotation.

### Analysis and visualization of non-canonical proteoforms

For the analysis of non-canonical proteoforms in **Figure 5**, DIAMOND v2.1.1.11.165 (Buchfink, Reuter, and Drost 2021) was used to perform BLASTp searches of the proteins against the UniProt database (release 2025_01) and BLASTp searches of the peptides against both UniProt and NCBI non-redundant (downloaded on 02/03/2025). Gene set enrichment analysis in **Figure 5C** was conducted using gget version 0.29.0 (Luebbert and Pachter 2023). Pfam (Mistry et al. 2021) searches were conducted using the pfam browser search tool: http://pfam.xfam.org/. Visualizations of ONT read coverage, peptides, ORFs, and transcript structure were done with our peptidescope package v0.0.1-beta.

### Code availability

Code for the ProteomeGenerator3 pipeline can be found here: https://github.com/kentsislab/proteomegenerator3. Code for peptidescope can be found here: https://github.com/shahcompbio/peptidescope. Code for sequence coverage calculators can be found here: https://github.com/shahcompbio/tcdo_pg_tools. The nf-core/nanoseq pipeline was implemented within the Isabl platform (Medina-Martínez et al. 2020).

### Reagents

Mass spectrometry grade (Optima liquid chromatography-mass spectrometry, LC−MS) water, acetonitrile (ACN), and methanol were purchased from Fisher Scientific (Fair Lawn, NJ). Formic acid (FA) and trifluoroacetic acid (TFA) of >99% purity were obtained from Thermo Scientific (Rockford, IL). All other reagents at MS-grade purity were obtained from Sigma-Aldrich (Saint Louis, MO).

### Proteome extraction and proteolysis

Cell pellets were resuspended in lysis buffer containing 8 M urea, 50 mM ammonium bicarbonate (pH 8.0), 5 mM CaCl_2_, and Halt protease and phosphatase inhibitor cocktail (Thermo Scientific). The pellet was lysed by sonication using S220 Covaris ultrasonicator device (at 175 W using 200 cycles per burst and a 10% duty factor at 4 °C for 6 min). The lysate was cleared by centrifugation at 14,000 × *g* for 15 min at 4 °C. The supernatant was then transferred to a new tube. Protein concentration was measured by Bradford Protein Assay Kit (Thermo Scientific). Proteins were reduced by addition of 10 mM dithiothreitol (DTT) and incubation for 45 min at RT. The cysteines were alkylated using 30 mM iodoacetamide for 45 min in dark at RT. For tryptic digestion, 1 mg of proteins were digested with trypsin (Promega) overnight at 37 °C with a 1:50 enzyme-to-protein ratio in 1 M urea. For LysC (Wako) digestion, a 1 mg protein aliquot was digested overnight at 37 °C with a 1:50 enzyme-to-protein ratio in 1 M urea. For LysN (ImmunoPrecise Antibodies) digestion, a 1 mg protein aliquot was digested overnight with a 1:50 enzyme-to-protein ratio in 1 M urea. For GluC (Promega) digestion, a 1 mg protein aliquot was digested overnight at 37 °C with a 1:50 enzyme-to-protein ratio in 0.5 M urea. For chymotrypsin (Promega) digestion, a 1 mg protein aliquot was digested overnight at 37 °C with a 1:100 enzyme-to-protein ratio in 1 M urea. For ArgC (Promega) digestion, a 1 mg protein aliquot was digested overnight at 37 °C with a 1:100 enzyme-to-protein ratio in 1 M urea. For AspN (Promega) digestion, a 1 mg protein aliquot was digested overnight at 37 °C with a 1:100 enzyme-to-protein ratio in 1 M urea. For Proalanase (Promega) digestion, a 1 mg protein aliquot was digested overnight at 37 °C with a 1:100 enzyme-to-protein ratio in 1 M urea, pH 1.5 adjusted with 37% HCl. The digestion was stopped by adding 100% TFA to the final concentration of 1%. The samples were cleared by centrifugation at 18,000 × *g* for 15 min at 4 °C and subsequently desalted on a 30-mg tC18 Sep-Pak cartridges (Waters).

### Peptide fractionation

Dried peptide pellets were resuspended in 5 mM KH₂PO₄/25% ACN, pH 2.7 at a concentration of approximately 1 µg/µL. Strong cation exchange (SCX) chromatography was performed using SCX macro-spin columns (Harvard Apparatus). Columns were first equilibrated with 5 mM KH₂PO₄/25% ACN before sample loading. Peptides were then applied to the column and centrifuged at 1,000 × g. Sequential elution was carried out using 300 µL of 5 mM KH₂PO₄/25% ACN containing increasing concentrations of KCl: 50 mM, 100 mM, 250 mM, and 500 mM. Eluates were collected in separate tubes. The resulting fractions were lyophilized prior to high-pH reversed-phase (RP) fractionation.

High-pH RP fractionation was performed using BioPureSPN Macro 100 mg Targa C18 columns (35–350 µg, HMM S18R, Harvard Apparatus/The Nest Group). The columns were first equilibrated with 100% methanol, followed by 0.1% TFA. Each SCX-peptide fraction, resuspended in 0.1% TFA, was loaded onto the column and washed with water. Peptides were eluted using an increasing gradient of ACN (7.5–60%) in 0.3% triethylamine (TEA) buffer (pH 10). In total, nine fractions were collected using: 7.5%, 10%, 12.5%, 15%, 17.5%, 20%, 22.5%, 25%, and 60% acetonitrile in 0.3% TEA. A final elution step was performed using 80% ACN/0.1% TFA. The samples were then concatenated into eight by combining as follows: 7.5% and 22.5%, 10% and 25%, 80% ACN/0.1% TFA and flow-through (FT). The resulting fractions were lyophilized before nano-liquid chromatography-ion mobility mass spectrometry (nLC-IM MS) analysis.

### LC-MS/MS analysis

Liquid chromatography–mass spectrometry (LC–MS) analysis was performed using a nanoElute 2 system (Bruker Switzerland AG) coupled to a timsTOF HT mass spectrometer (Bruker Daltonics GmbH & Co.). Peptide samples (∼200 ng in 1 µL) were directly loaded onto a 25 cm C18 reverse-phase column (75 µm inner diameter, 1.7 µm particle size; Aurora Series Ultimate, IonOpticks). Chromatographic separation was carried out at a flow rate of 400 nL/min using a linear gradient from 2% to 35% buffer B (99.9% acetonitrile, 0.1% formic acid) over 45 minutes, followed by an increase to 95% buffer B sustained for 8 minutes (total run time: 53 minutes). The column temperature was maintained at 50 °C using an integrated column heater. Data-independent acquisition using parallel accumulation–serial fragmentation (dia-PASEF) was employed, utilizing 32 isolation windows. Each window had a width of 26 Da with a 1 Da overlap and was configured with variable ion mobility (IM) settings. Two acquisition steps were used, yielding a total cycle time of 1.80 seconds. The m/z range was set from 400 to 1201, and the ion mobility range (1/K₀) spanned from 0.60 to 1.60 V·s/cm². Ramp and accumulation times were both set to 100 ms. Collision energy was dynamically adjusted from 20 to 59 eV across the defined ion mobility range.

### Proteomics data analysis with Fragpipe

diaPASEF data acquired on the timsTOF HT instrument were analyzed using FragPipe (version 22.0) with the “DIA_SpecLib_Quant_diaPASEF” workflow to generate a spectral library and perform quantification for the raw .d files. Within diaTracer, “Delta Apex IM” was set to 0.01, “Delta Apex RT” was set to 3, “RF max” was set to 500, and “Corr threshold” was set to 0.3. The mass defect filter was enabled. In the MSFragger database search, the initial precursor and fragment mass tolerances were set to 10 ppm and 20 ppm, respectively. Spectrum deisotoping, mass calibration, and parameter optimization were enabled. The isotope error was set to “0/1/2”. The A673 protein search database (i.e., the protein fasta output generated by PG3), appended with common contaminants and decoys, was used in the search. Enzyme specificity was set to “stricttrypsin” and the maximum allowed missed cleavages were set to 2. Oxidation of methionine and N-terminal acetylation were set as variable modifications. Carbamidomethylation of cysteine was used as a fixed modification. The maximum number of variable modifications for each peptide was set to 3. MSBooster and Percolator were used to predict the RT and MS/MS spectra, and to rescore PSMs. The final FDR-filtered PSMs and the pseudo-MS/MS mzML spectral files were used by EasyPQP to generate the spectral library. The spectral library was then used with the DIA-NN (v1.8.2) quantification module to quantify the library peptide ions in the individual diaPASEF data. The same settings were employed for all proteolytic enzymes.

To conduct the search against the concatenated A673-specific proteome and SwissProt human proteome, we first downloaded the canonical + isoforms human SwissProt protein database (UniProt, downloaded September 6^th^, 2024), and appended with common contaminants and decoy sequences. We then concatenated the A673-specific proteome used in the previous searches (which we had already appended with common contaminants and decoy sequences) with this SwissProt fasta and used seqkit (v2.10.0) to remove any duplicate sequences. We then used the same search parameters described in the previous paragraph to search this protein fasta file.

### Proteomics data analysis with DIA-NN

In the analysis of timsTOF HT diaPASEF tryptic data, search parameters of DIA-NN (version 2.1.0) were set as follows: precursor FDR 1%; mass accuracy at MS1 and MS2 set to 15 ppm as recommended; scan window set to 6 as recommended; protein inference and MBR turned on; scoring set to generic; proteotypicity at gene level; quantification strategy set to Quant UMS (high precision); machine learning set to NNs (cross-validated); cross-run normalization set as RT-dependent. The search was performed in the library-free mode. The settings for in silico library generation as follows: Trypsin/P with maximum 1 missed cleavage; protein N-terminal M excision on; carbamidomethyl on C as fixed modification; no variable modification; peptide length from 7 to 30; precursor charge 1–4; precursor m/z from 300 to 1800; fragment m/z from 200 to 1800.

### Proteomics data analysis with Spectronaut

In the directDIA+ (Deep) analysis of timsTOF HT diaPASEF tryptic data, search parameters of Spectronaut (version 19.0) were set as follows: enzyme specificity was set to “Trypsin/P”; digest type was set to specific with maximum 2 missed cleavages; peptide length from 7 to 52; oxidation of methionine and N-terminal acetylation were set as variable modifications; carbamidomethylation of cysteine was used as a fixed modification; precursor PEP cutoff 0.2; precursor *q* value cutoff 0.01; protein *q* value cutoff 0.01 at experiment level and 0.05 at run level; protein PEP cutoff 0.75. Other parameters were set as default. Peptide (MS1) and precursor ion (MS2) extracted ion chromatogram (XIC) profiles for the EWSR1:FLI1 fusion peptide GQQNPSYDSVRRGAW.3 (shown in **Supplementary Figure 5**) were generated using Spectronaut (version 19.0), applying parameters consistent with those described above, except that the enzyme specificity was set to Chymotrypsin/P.

### Enhancing DIA-PASEF Coverage for Non-Canonical Proteolytic Digests

Non-fractionated chymotryptic peptide samples were used to evaluate various DDA- and DIA-PASEF acquisition methods aimed at addressing non-tryptic peptide diversity in DIA workflows (also see section *Proteome extraction and proteolysis*). Liquid chromatography–mass spectrometry (LC–MS) analysis was performed using a nanoElute 2 system (Bruker Switzerland AG) coupled to a timsTOF Ultra 2 mass spectrometer (Bruker Daltonics GmbH & Co.). Peptide samples (∼100 ng in 1 µL) were directly loaded onto a 25 cm C18 reverse-phase column (75 µm inner diameter, 1.5 µm particle size; PepSep Ultra, Bruker Daltonics GmbH & Co.). Chromatographic separation was carried out at a flow rate of 300 nL/min using a linear gradient from 2% to 22% buffer B (99.9% acetonitrile, 0.1% formic acid) over 25 minutes, then an increase to 30% buffer B over 5min, followed by an increase to 95% buffer B sustained for 8 minutes (total run time: 38 minutes). The column temperature was maintained at 50 °C using an integrated column heater. The timsTOF Ultra DDA- and DIA-PASEF MS-acquisition methods are summarized in **Supplementary table 2.** Standard and Double Rainbow DIA-PASEF data were analyzed using FragPipe (version 23.0) with the ‘DIA_SpecLib_Quant_diaPASEF’ workflow, while all DDA-PASEF data files were processed using the ‘LFQ-MBR’ workflow. Searches were conducted against the A673-specific protein database generated by PG3, appended with common contaminants and decoy sequences, as described previously.

## Acknowledgements

We thank the MSK High Performance Computing Group, Proteomics Core, and Integrated Genomics Operation Core for technical support and members of our laboratories for useful discussions. Additionally, we thank Jeffery Summers, Wenming Xiao, Devaveena Dey, Alexey I. Nesvizhskii, Fengchao Yu, and the Nesvizhskii Lab for their helpful suggestions. This work was supported by the FDA U01 FD008299, NIH R01 CA204396 and P30 CA08748, Break Through Cancer Foundation, Starr Cancer Consortium, Alan B. Slifka Foundation, and Pediatric Cancer Foundation. AK is a Scholar of the Leukemia & Lymphoma Society. Figures were generated using BioRender, with individual figure citations provided per license (*BioRender.com*).

## Supplement

**Supplementary Figure 1.**
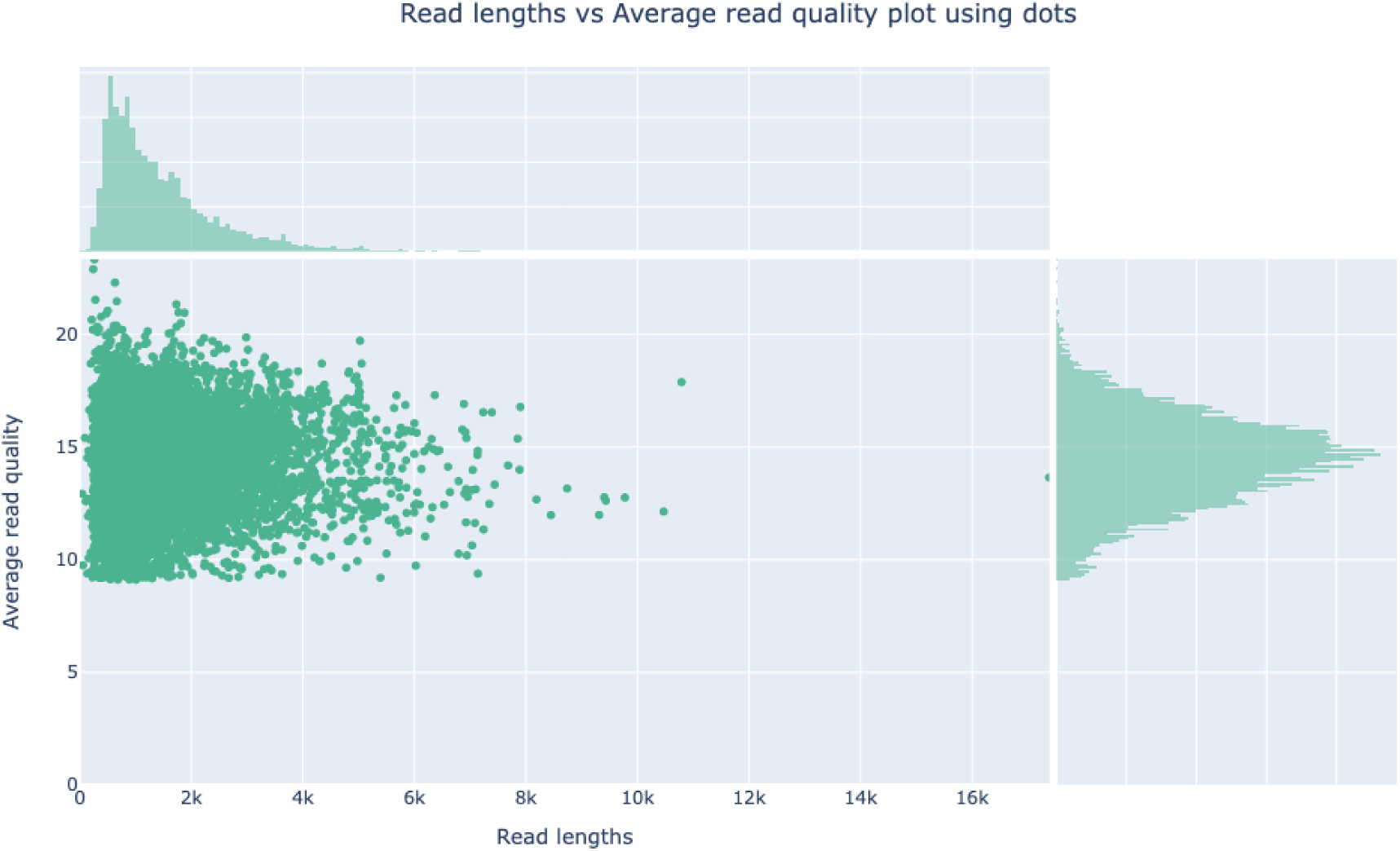
Raw read distributions from long-read cDNA sequencing of the A673 cell line from Nanoplot. Y-axis of main plot shows Phred scores and x-axis shows read lengths in kilobases.

**Supplementary Figure 2.**
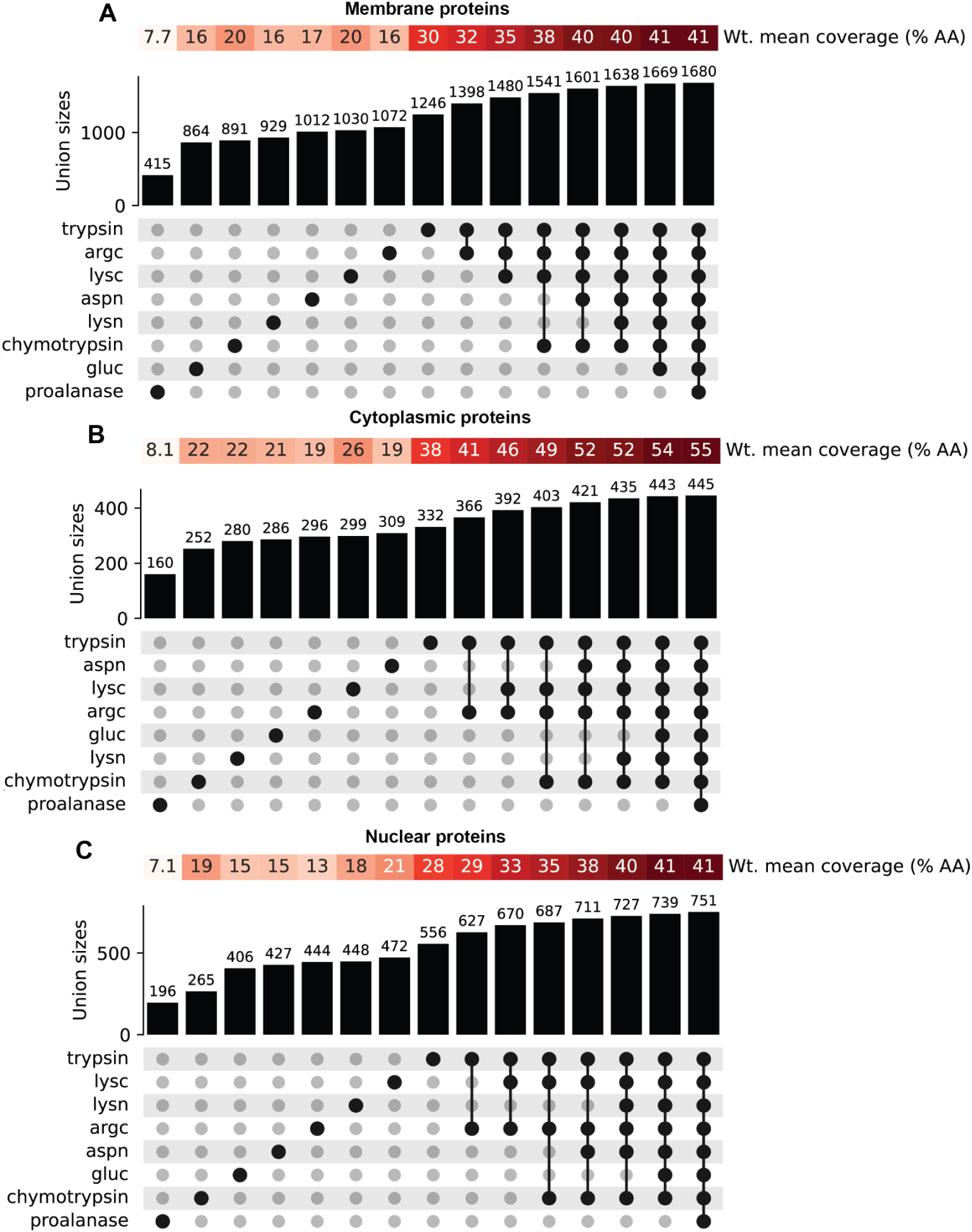
Bar plots highlighting the total number of protein IDs identified with different enzyme combinations for membrane (*A*), cytoplasmic (*B*), and nuclear (*C*) proteins. Corresponding mean amino acid sequence coverage is shown above each bar. Matrix details which enzyme is used in the combination.

**Supplementary Figure 3.**
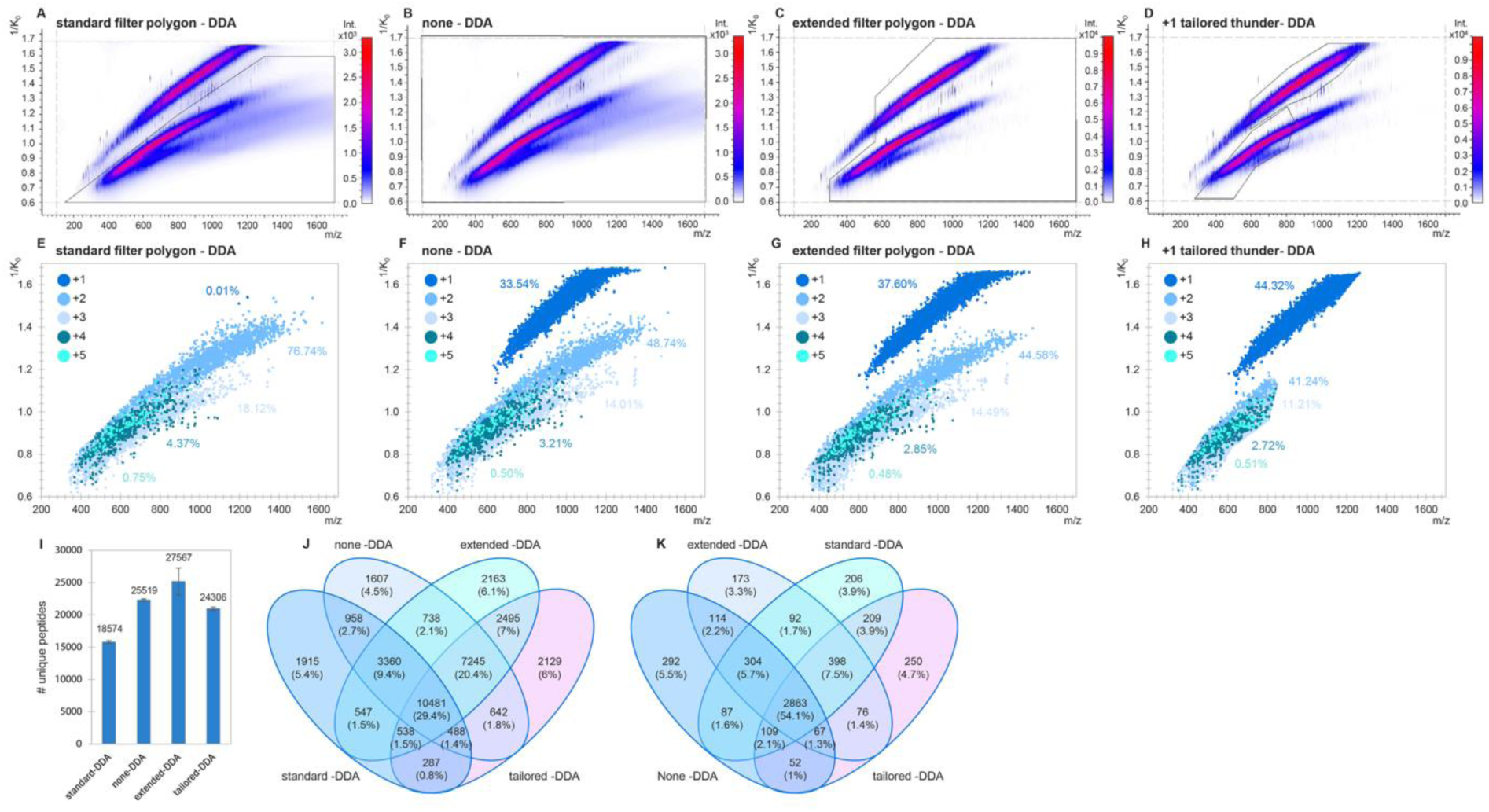
Evaluation of different DDA-PASEF fragmentation isolation polygons for the identification of singly (+1) and multiply charged (≥+2) chymotryptic peptides. A-D) Exemplary heat maps of ion intensities (blue-red scale) across the inversed ion mobility (1/K_0_) vs m/z dimensions showing fragmentation events (black line). **E-H)** All peptides identified across the 1/K_0_ vs m/z dimensions colored by charge state. The percentages of detected singly charged (+1) and multiply charged (≥+2) peptides are also highlighted for the none-, extended-, standard-, and +1 tailored-DDA-polygon acquisition strategies. **I)** Average number of unique peptides identified per injection for each method (error bars represent *mean±stdev*, n = 3). Venn diagram s illustrating the overlap in identified **J)** unique peptides and **K)** proteins among the four tested DDA-PASEF isolation window strategies.

**Supplementary Figure 4.**
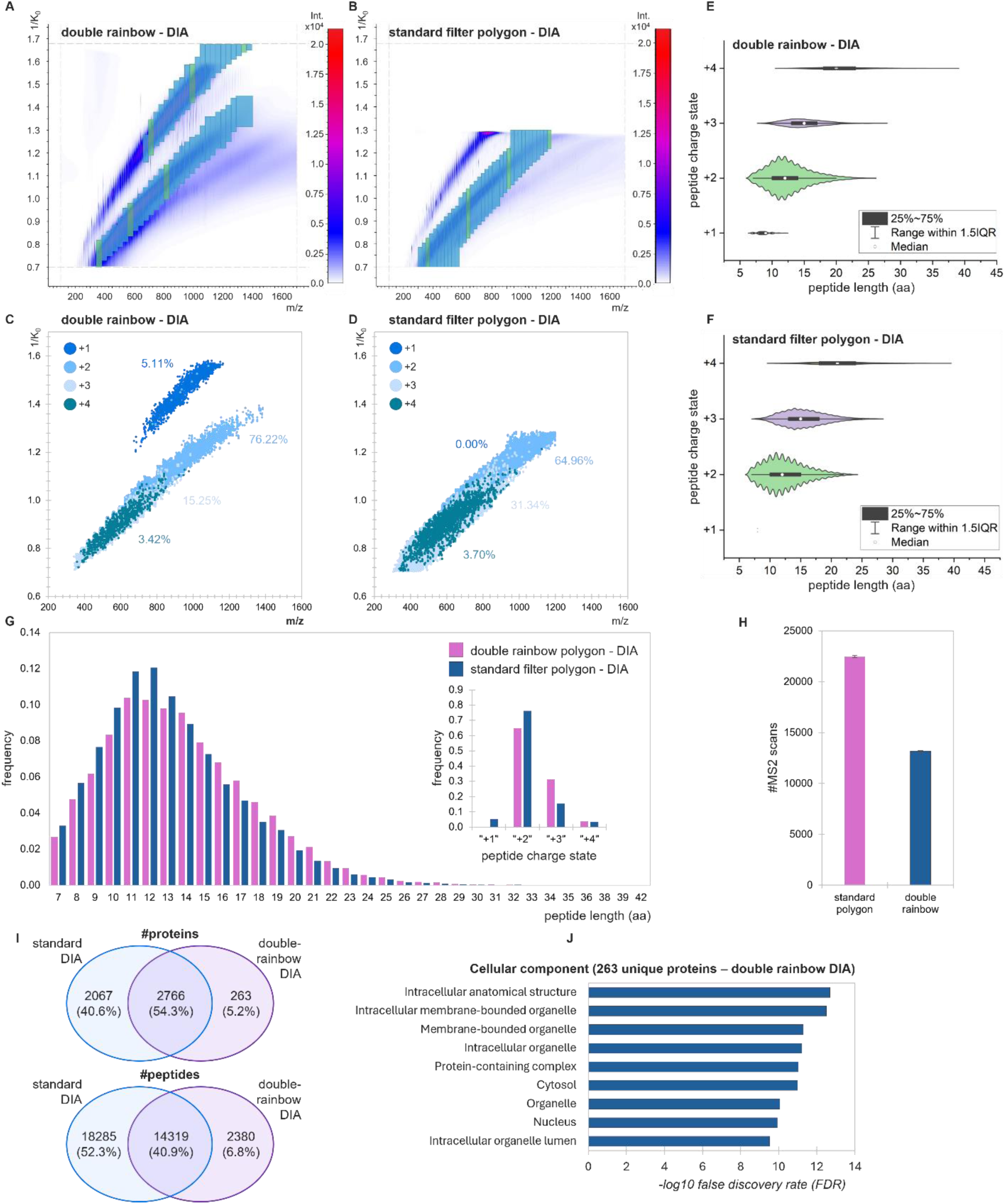
Evaluation of different DIA-PASEF fragmentation isolation window strategies for the identification of singly charged (+1) and multiply charged (≥+2) chymotryptic peptides. A–B) Representative heatmaps of ion intensities (blue-to-red scale) across the inverse ion mobility (1/K₀) vs. m/z dimensions, indicating fragmentation events (blue boxes, single injection). **C–D)** Distribution of all identified peptides across the 1/K₀ vs. m/z dimensions, color-coded by charge state. The percentages of detected singly charged (+1) and multiply charged (≥+2) peptides are highlighted for the standard and double-rainbow DIA-PASEF acquisition strategies. **E–F)** Peptide length and charge state distributions (7–50 amino acids) for the standard and double-rainbow DIA-PASEF isolation windows. The total number of identified peptides for each specific charge state is indicated in blue. **G)** Frequency distribution of identified peptide lengths and charge states using the Standard and Double Rainbow DIA-PASEF approaches. **H)** Number of MS2 scans triggered per injection in each method (error bars represent mean±stdev, n = 3). **I)** Venn diagrams comparing the numbers of identified proteins and peptides between the standard and double-rainbow DIA-PASEF isolation window strategies. **J)** Gene Ontology (GO) cellular component analysis performed on 263 uniquely identified proteins obtained using the Double Rainbow DIA-PASEF approach. Enrichment analysis was conducted using the STRING v12.0 database^92^.

**Supplementary Figure 5.**
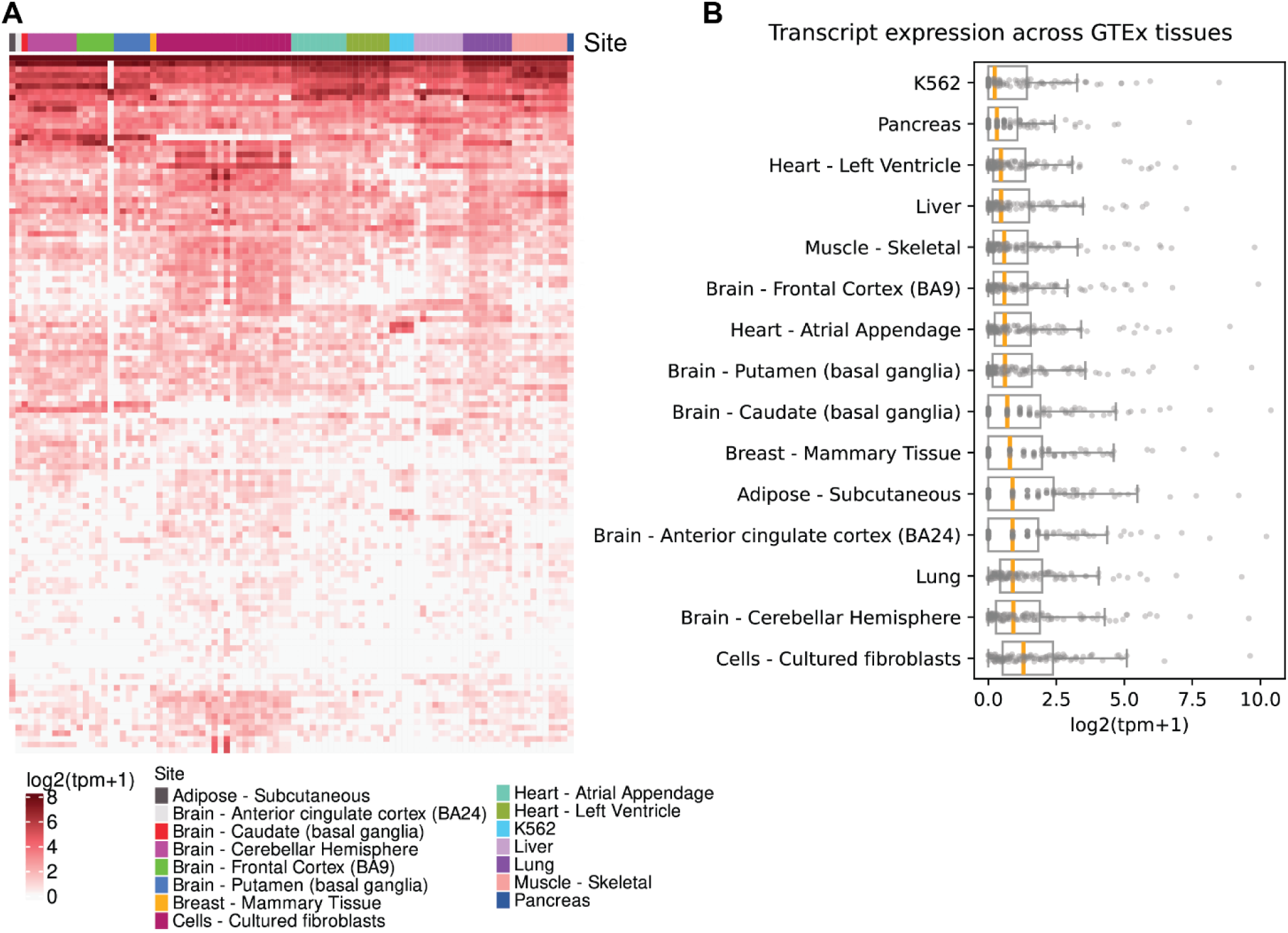
Expression of non-canonical transcripts in the A673 transcript assembly which overlapped with the GTEx annotation. (**A**) Heatmap of log-transformed transcripts per million (TPM) for individual transcripts, in which each column represents a different GTEx sample, and each row represents a different transcript. (**B**) Tukey boxplots with strip plots overlaid depicting log-transformed TPMs, in which each point represents the mean log2(TPM+1) for a given transcript taken over all samples in a given site.

**Supplemental Figure 6.**
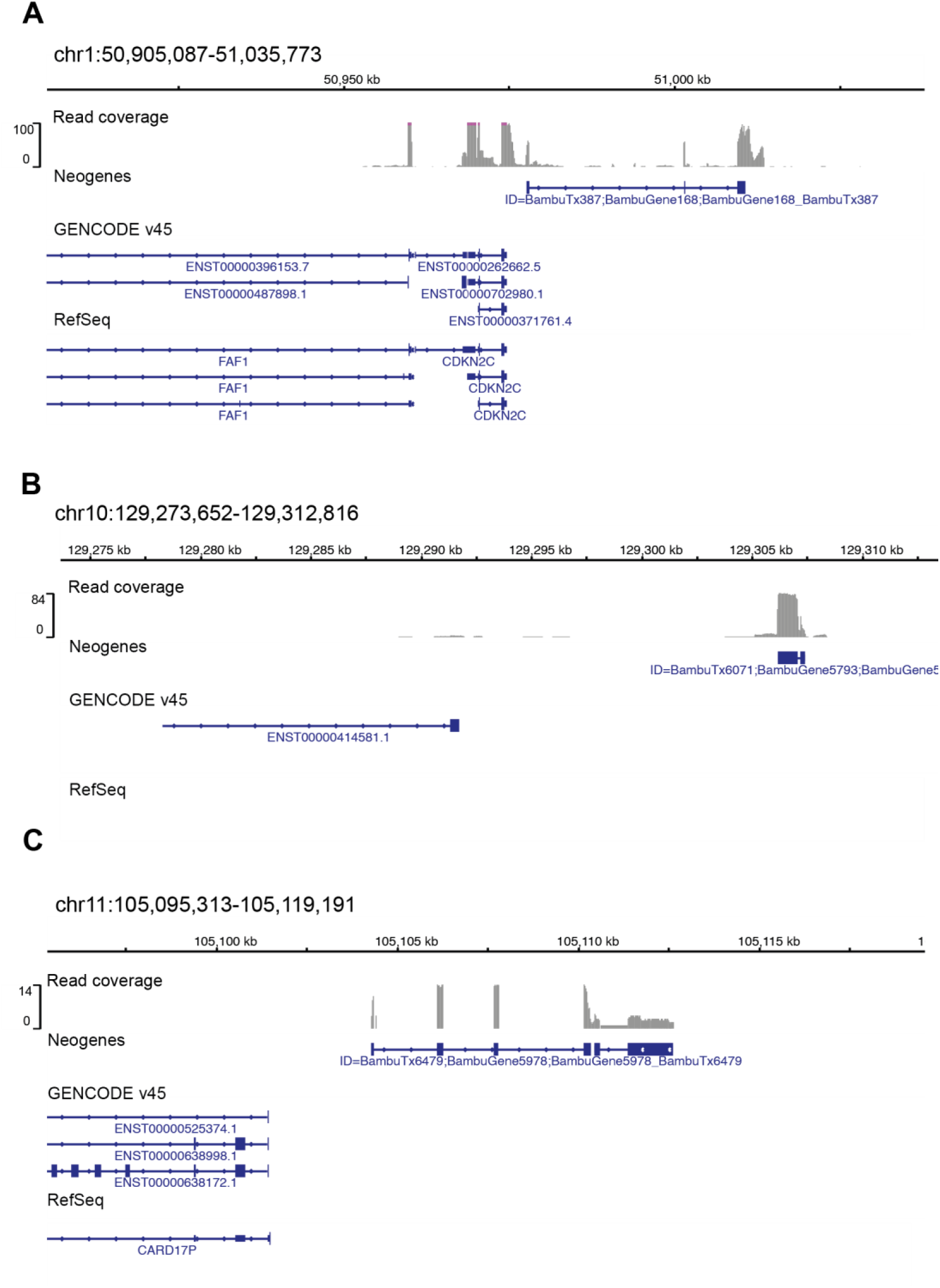
Examples of A673 neogenes appearing in gene deserts (previously unannotated genomic regions). For each panel, tracks from top to bottom show: ONT read coverage, transcript structure of detected neogenes, GENCODE v45 and RefSeq (downldetected peptides (orange) shown on transcript structure (green), detected transcripts shown with predicted ORFs (the unknown protein is highlighted in green), and RefSeq (downloaded from UCSC on 07/09/2025).

**Supplementary Figure 7.**
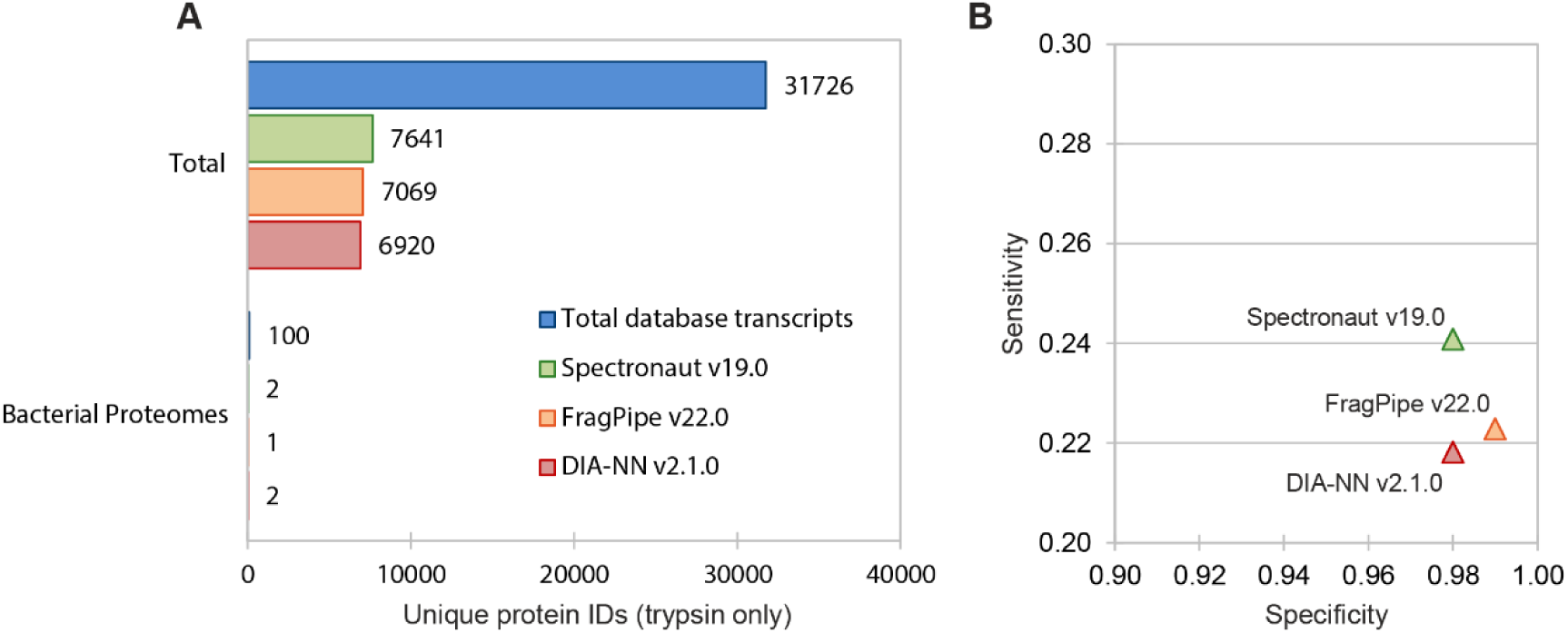
Benchmarking of computational proteomics search methods. **A)** Bar plot depicts the number of quantified proteoforms in the A673 tryptic dataset analyzed using Spectronaut 19 (direct DIA), FragPipe+MSFragger v22.0 (diaTracer workflow), and DIA-NN v1.9.1 (library-free mode). The A673 PG3 protein sequence database, including nonhuman *Archaebacteria loki* proteins, was used for analysis. **B)** Sensitive vs specificity for different computational proteomics search methods using the results of panel *A*. Sensitivity was calculated as the fraction of identified protein groups relative to the total number of annotated A673 transcripts. Specificity was assessed as the inverse proportion of erroneously matched *A. loki* sequences among all identifications.

**Supplementary Figure 8.**
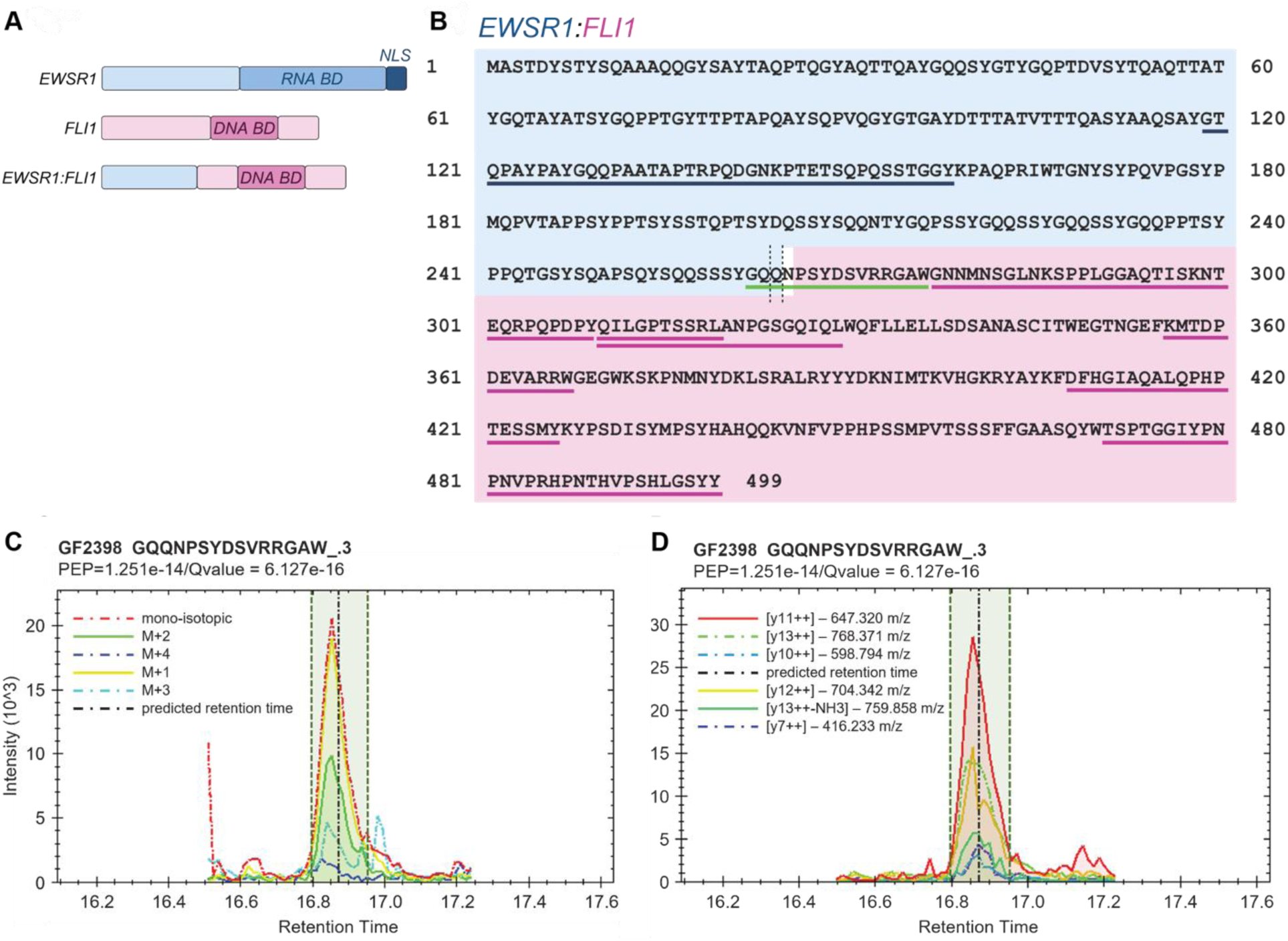
EWSR1:FLI1 fusion protein amino acid sequence and MS-identified chymotryptic peptides. **A)** Structure of EWS, FLI1, and EWSR1:FLI1. *Abbr. BD – Binding Domain, NLS – Nuclear Localization Signal/Sequence*. **B)** The full-length amino acid (aa) sequence of the EWSR1:FLI1 fusion protein is shown, encompassing the N-terminal domain derived from EWSR1 (in blue) and the C-terminal ETS DNA-binding domain contributed by FLI1 (in purple). The breakpoint resulting from the chromosomal translocation t(11;22)(q24;q12) in A673 is indicated by vertical lines separating the EWSR1 and FLI1 segments. Peptides identified by LC-MS/MS are underlined, demonstrating sequence coverage across the fusion junction as well as within both EWSR1- and FLI1-derived regions (34%). The presence of peptide spanning the fusion breakpoint (underlined in green) confirms expression of the chimeric protein at the proteomic level. Peptide (MS1) **C)** and precursor ion (MS2) eXtracted Ion Chromatogram (XIC) **D)** profiles for EWSR1:FLI1 fusion protein with precursor GQQNPSYDSVRRGAW.3. *Panel A and B created with BioRender.com;* https://BioRender.com/vgy5ec2.

**Supplementary Figure 9.**
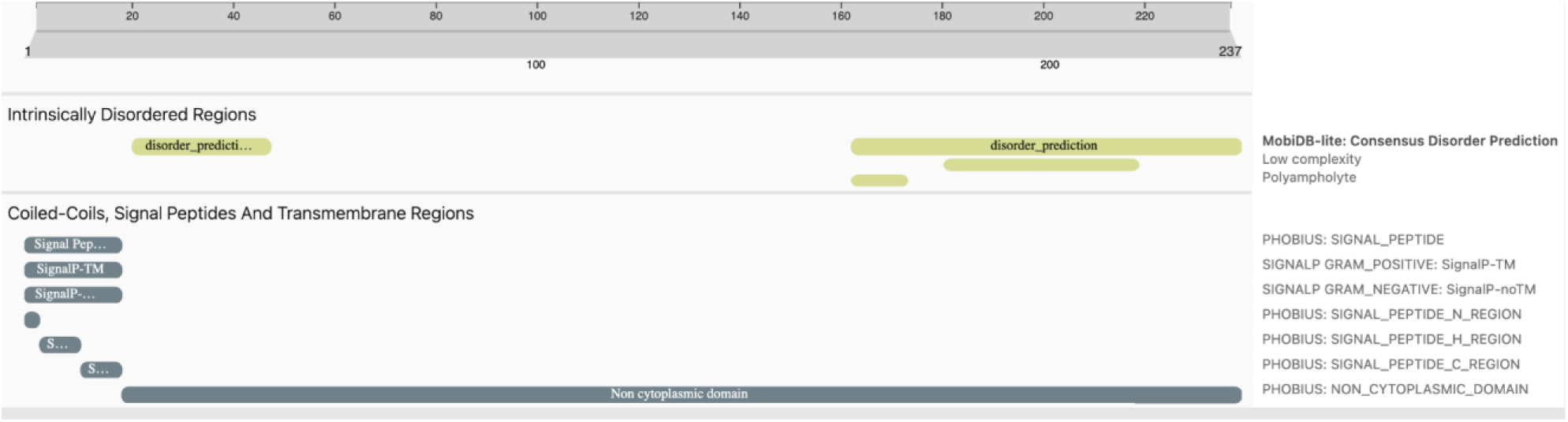
Pfam search results for non-canonical EXOC7 proteoform. Pfam search results corresponding to the non-canonical proteoform depicted in Figure 5E. For this proteoform, we found a unique tryptic peptide (PEP = 2e-4) related to an alternative ORF on ENST00000465252.5. 12 pfam domain matches were found (shown above).

**Supplementary Table 1.**
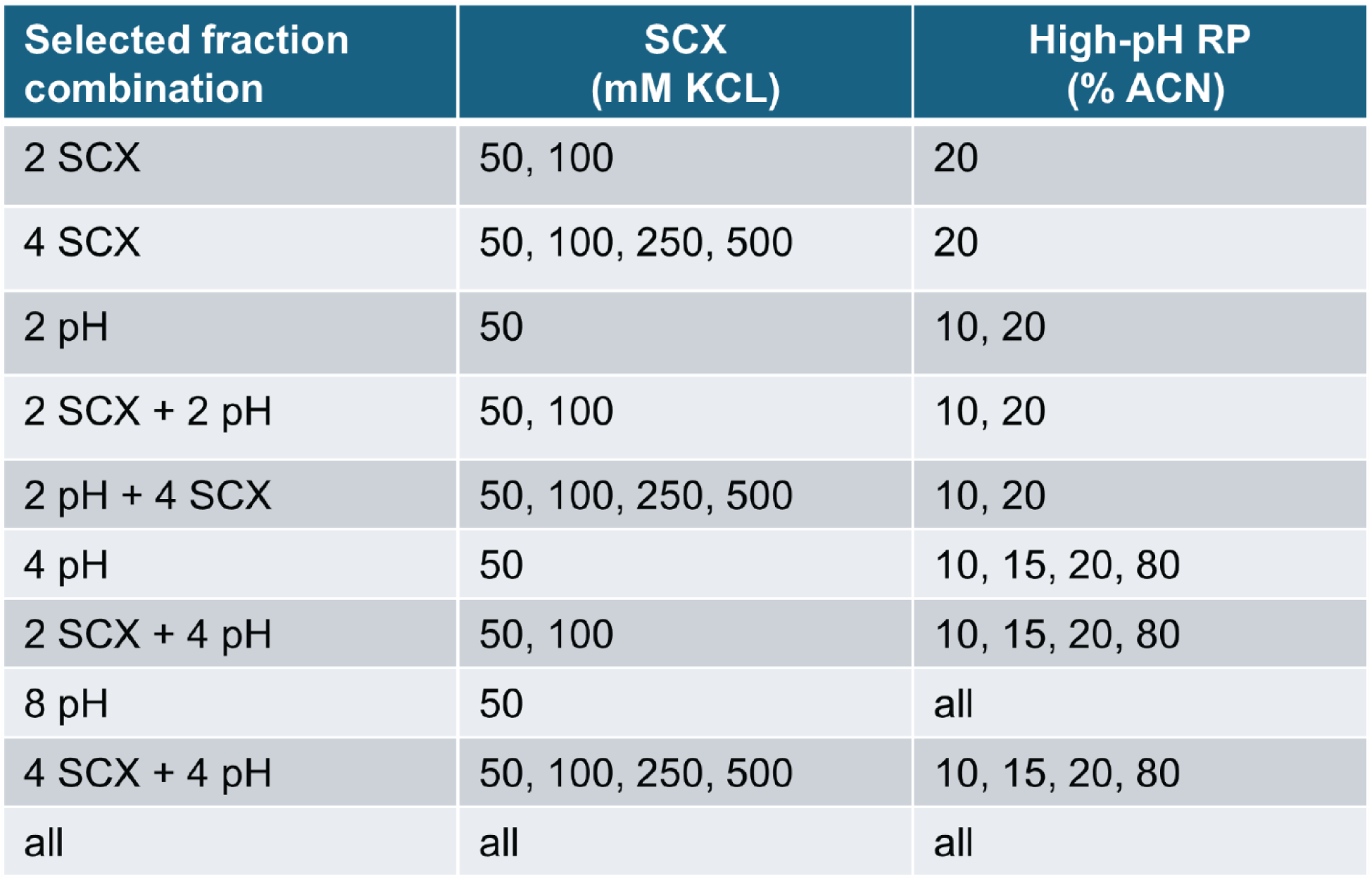
Evaluation of SCX and High-pH RP fractionation strategies to optimize fraction numbers without compromising proteome coverage.

| Selected fraction combination | SCX (mM KCL) | High-pH RP (% ACN) |
| --- | --- | --- |
| 2 SCX | 50, 100 | 20 |
| 4 SCX | 50, 100, 250, 500 | 20 |
| 2 pH | 50 | 10, 20 |
| 2 SCX + 2 pH | 50, 100 | 10, 20 |
| 2 pH + 4 SCX | 50, 100, 250, 500 | 10, 20 |
| 4 pH | 50 | 10, 15, 20, 80 |
| 2 SCX + 4 pH | 50, 100 | 10, 15, 20, 80 |
| 8 pH | 50 | all |
| 4 SCX + 4 pH | 50, 100, 250, 500 | 10, 15, 20, 80 |
| all | all | all |

**Supplementary Table 2.** Summary of MS acquisition method settings used for each optimized DDA- and DIA-PASEF isolation polygon. The accompanying Excel file includes detailed method parameters, including:

- MS_DDA: Optimized DDA-PASEF acquisition methods tailored to improve the detection of non-tryptic peptides in data-dependent workflows.
- DDA_Isolation_Polygon_Coordinates: Coordinates defining the boundaries of the tested DDA isolation polygons used for precursor ion selection and fragmentation.
- MS_DIA: Optimized DIA-PASEF acquisition methods designed to enhance coverage of non-tryptic peptides in data-independent workflows.
- DIA-PASEF_Window_Settings: Coordinates defining the m/z and ion mobility boundaries of the isolation windows used for precursor ion selection and fragmentation in the standard- and double-rainbow DIA-PASEF acquisition schemes.

**Supplementary Table 3.** Table of the 72 putative non-canonical proteoforms which had no match when DIAMOND BLASTp was run against UniProt. Each row contains information for a peptide for one of the detected proteins, including the enzyme which detected the peptide.

**Supplementary Table 4.** Table of the 17 putative non-canonical proteoforms which had no match in DIAMOND BLASTp searches against UniProt and were detected in a computational proteomics search against a protein search database composed of the A673-specific proteome and the SwissProt human reference proteome (a subset of Supplementary Table 3). Rows are the same as Supplementary Table 3.

